# Open-ended molecular recording of sequential cellular events into DNA

**DOI:** 10.1101/2021.11.05.467507

**Authors:** Theresa B. Loveless, Courtney K. Carlson, Catalina A. Dentzel Helmy, Vincent J. Hu, Sara K. Ross, Matt C. Demelo, Ali Murtaza, Guohao Liang, Michelle Ficht, Arushi Singhai, Marcello J. Pajoh-Casco, Chang C. Liu

## Abstract

Genetically encoded DNA recorders noninvasively convert transient biological events into durable mutations in a cell’s genome, allowing for the later reconstruction of cellular experiences using high-throughput DNA sequencing^1^. Existing DNA recorders have achieved high-information recording^2–15^, durable recording^3,5–10,13,15–19^, multiplexed recording of several cellular signals^5–8,19,20^, and temporally resolved signal recording^5–8,19,20^, but not all at the same time in mammalian cells. We present a DNA recorder called peCHYRON (prime editing^21^ Cell HistorY Recording by Ordered iNsertion) that does. In peCHYRON, mammalian cells are engineered to express prime editor and a collection of prime editing guide RNAs^21^ (pegRNAs) that facilitate iterative rounds of prime editing. In each round of editing, prime editor inserts a variable triplet DNA sequence alongside a constant propagator sequence that deactivates the previous and activates the next step of insertion. Editing can continue indefinitely because each insertion adds the complete sequence needed to initiate the next step. Because only one active target site is present at any given time, insertions accumulate sequentially, in a unidirectional order. Thus, temporal information is preserved in the order of insertions. Durability is achieved through the use of a prime editor that only nicks a single DNA strand, effectively avoiding deletion mutations that could otherwise corrupt the information stored at the recording locus. High-information content is established by co-expressing a variety of pegRNAs, each harboring unique triplet DNA sequences. We demonstrate that constitutive expression of such a library of pegRNAs generates insertion patterns that support straightforward reconstruction of cell lineage relationships. In an alternative pegRNA expression scheme, we also achieve multiplexed recording by manually pulsing expression of different pegRNAs, then reconstructing pulse sequences from the peCHYRON records. Additionally, we coupled the expression of specific pegRNAs to specific biological stimuli, which allowed temporally resolved, multiplexed recording of chemical exposures in populations of mammalian cells.

**Graphical abstract:** 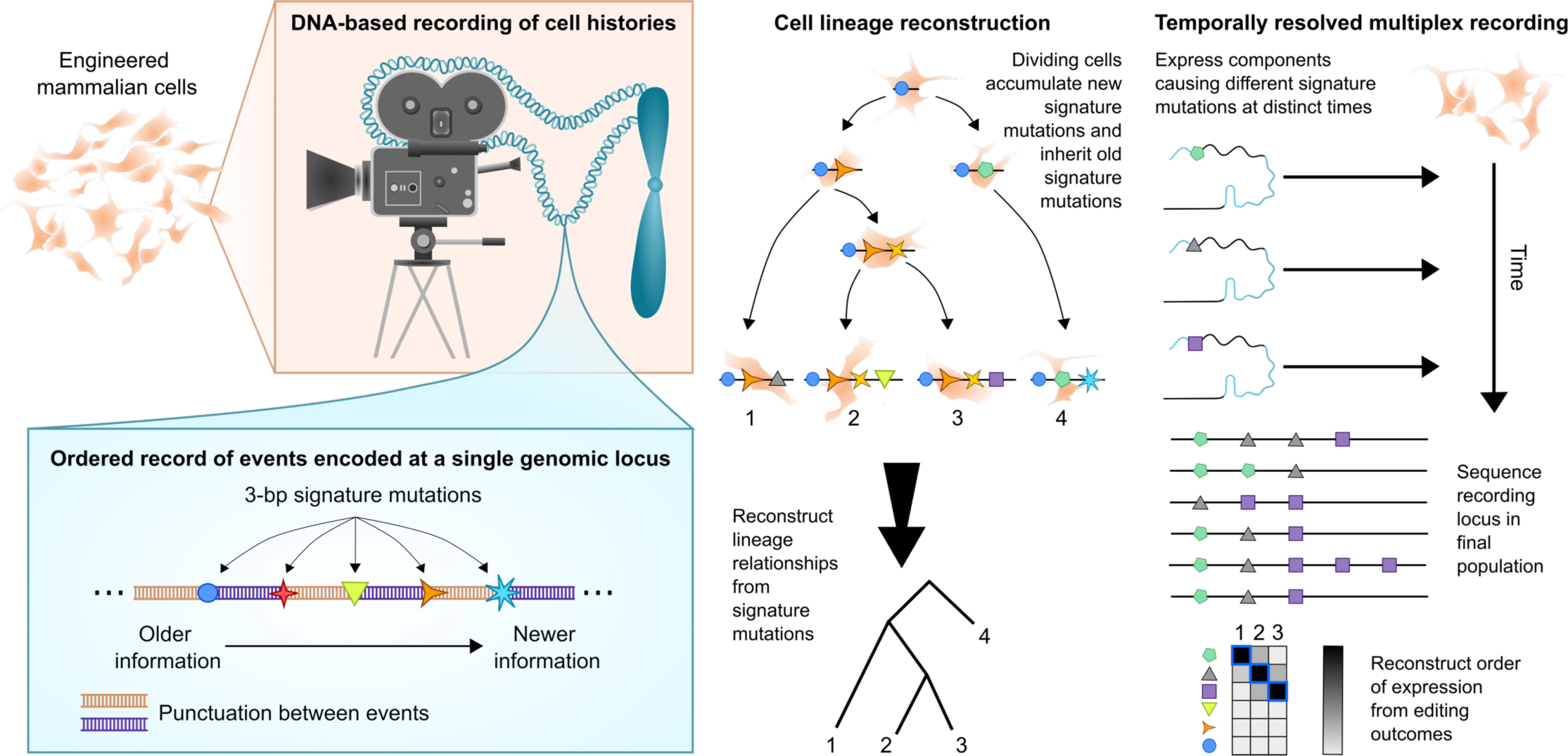

---

Understanding and eventually engineering development will require scientists to relate both cell lineage and (sequential) patterns of signaling to specific cell fates. To achieve this goal, it will be necessary to track the phylogenies of individual cells, and also to follow many signaling pathways in parallel over long timescales at single cell resolution as those cells differentiate. DNA recording can make these analyses possible by enabling each cell to record transient phenomena (*e.g.,* the existence of an ancestral stem cell that undergoes apoptosis after giving rise to differentiated daughter cells, or the presence of fleeting microenvironmental cues that affect gene expression within the cell of interest) and permanently store a record of the transient event as a mutation in its DNA (**Figure 1a**). To maximize the amount of information that can be extracted from its records, a DNA recorder should have the following properties: (1) high information content: the ability to generate records with diverse sequences, either random sequences that can be used to uniquely barcode individual cells and therefore mark distinct cell lineages with high confidence, or sequences that encode patterns of transient events; (2) order: the ability to record events in the sequence they arrive, so that the timing of events can be deciphered from the relative positions of mutations in the records; (3) punctuation: the ability to delimit one event from the next in the record; (4) durability: the ability to record new events without corrupting or deleting the record of older events; (5) continuous operation: the ability to record new events over long enough timescales to study biologically interesting processes; and (6) multiplexability: the ability to record multiple distinct signals in parallel.

**Figure 1.**
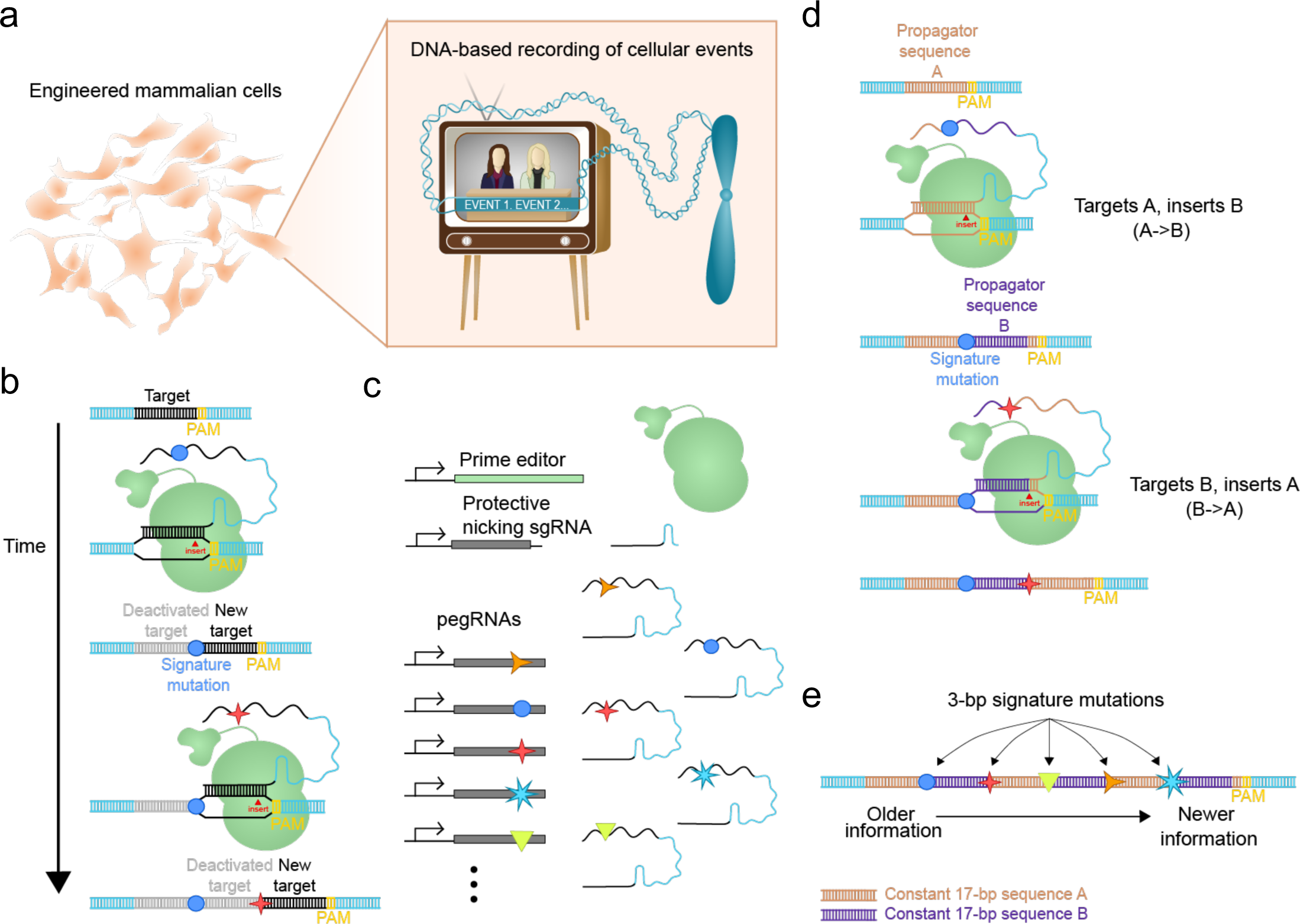
Overview of peCHYRON. **a**, Using peCHYRON, transient events are recorded into DNA in the order they occur, as in a scrolling chyron. **b**, Iterative sequential insertion of signature mutations via prime editing. Signature mutations are variable 3-bp sequences, each theoretically encoding up to 6 bits of information. In a single cycle of prime editing, a signature mutation and a 17-bp propagator sequence are inserted at the recording locus adjacent to the PAM. The newly-inserted propagator sequence forms the target site for the next cycle of editing because it is now PAM-adjacent. The previous target site is deactivated due to separation from the PAM. This process can repeat indefinitely, resulting in ordered accumulation of regularly-spaced signature mutations. **c**, A collection of pegRNAs facilitates the insertion of signature mutations at the recording locus during continuous and iterative editing. **d**, Propagation with alternating pegRNAs. To avoid issues with repetitive sequence elements, the recording locus alternates between addition of two different 17-bp propagator sequences. In one step, a pegRNA targets sequence A and inserts sequence B. In the next step, a pegRNA targets sequence B and inserts sequence A. The process repeats and continuous propagation of the recording locus can proceed without end. **e**, How to interpret information at a peCHYRON locus. Information-rich signature mutations are separated by constant 17-bp propagator sequences, resulting in clear punctuation between sequential editing events. The age of a signature is reflected by its position in the record, with PAM-proximal signatures reporting on the most recent events.

Before the invention of prime editor^21^, DNA recorders suffered from fundamental mechanistic tradeoffs that prevented them from achieving all these properties simultaneously^2–14,16–19^. Legacy recorders either caused double-strand breaks in DNA, compromising durability because corruptive mutations would arise during DNA repair, or could install only a limited number of potential mutations (*e.g.*, base editor recorders). However, DNA recorders that use prime editor, a nickase Cas9-reverse transcriptase (RT) fusion that installs precise mutations specified by pegRNAs^15,22^, do not suffer from these same limitations^15,22^. For example, DNA Typewriter^15,22^, which uses prime editor to make small insertion mutations in temporal order on a synthetic recording locus, can be used to resolve the order of 16 transfections of pegRNA-encoding plasmids or retrace the full lineage relationships between thousands of cells. peCHYRON is an alternative approach. As in DNA Typewriter, prime editor is used to make ordered, high-information, non-corruptive mutations. Random mutations can be used to retrace cell lineage, or particular mutations can be tied to particular events. Unlike DNA Typewriter, peCHYRON adds a new target site in each round of recording, meaning that peCHYRON recording does not require a synthetic locus to record at and can, in principle, be extended for an arbitrarily high number of rounds (**Figure 1b**).

## Architecture of peCHYRON

Our goal with peCHYRON was to build a recorder that couples high durability and extensibility with the high information content that is easily contained in mutations installed by prime editor. To achieve high durability, we used the PE2 instantiation of prime editor, in which only one DNA strand is nicked^21^, so that the unintended mutation rate is extremely low (∼1% as frequent as the intended edit). To achieve high extensibility, we designed peCHYRON so that each round of recording installs a target site for the next round. First, a 20-bp locus in a mammalian cell’s genome is targeted by a pegRNA. The pegRNA directs prime editor (PE2max)^21,23^ to reverse transcribe the pegRNA’s programmable template sequence into the locus. The reverse transcriptase (RT) template sequence is programmed to insert a variable 3-nt sequence that encodes up to 6 bits of information followed by a constant 17-nt propagator sequence that serves as the target sequence for the next insertion step (the 3 bp nearest to the PAM are not altered during an edit, so only 17 nts of the target need to be re-installed to constitute a new 20-bp target adjacent to the PAM). After one cycle of insertion, the previous propagator sequence is no longer PAM-adjacent, and therefore inactive, while the new propagator sequence that forms the new 20-bp target site becomes PAM-adjacent, and therefore active. The next step of editing inserts another variable 3-nt sequence along with the next propagator sequence, iterating indefinitely. In this manner, each variable 3-bp sequence, which we refer to as a signature mutation, records an event. Because the information content of the signature mutation can be up to 6 bits, peCHYRON has sufficient complexity to theoretically record 64 different types of events – each different signature is installed by its own pegRNA (**Figure 1c**), which could be expressed inducibly. New signature mutations appear in the order of events recorded. Constant 17-bp sequences separate signature mutations, acting as punctuation. The recording process does not change previous records, as edits occur through sequential insertions, and recording can, in principle, propagate continuously without end (**Figure 1b**).

The simplest implementation of peCHYRON would involve the sequential insertion of a constant propagator sequence (**Extended Data Figure 1a-c**). However, iterative addition of a single propagator sequence requires a pegRNA whose RT template sequence (containing the 17-nt propagator sequence to be inserted) and primer binding sequence (PBS)^21^ share identity. During the RT step of prime editing, this would allow primer binding in the pegRNA’s RT template sequence rather than the PBS, resulting in no net insertion (**Extended Data Figure 1c**). We therefore settled on the use of two alternating 17-bp propagator sequences in our peCHYRON design so that the RT template sequence and PBS in each pegRNA are distinct. The peCHYRON architecture we implemented is shown in **Figure 1d** where one pegRNA targets site A and adds a propagator sequence B (A→B) and the other pegRNA targets sequence B and adds a propagator sequence A (B→A), allowing the cycle to continue. Importantly, although peCHYRON uses alternating propagator sequences, this design does not require 2 edits to record a single event. For any event that we wish to record, we simply express a pair of A®B and B®A pegRNAs simultaneously, so one of the pegRNAs corresponding to the event is compatible with editing the recording locus at any time. Recorded information in the resulting peCHYRON locus is easily interpretable (**Figure 1e**).

## Identification of efficient parts for peCHYRON

In the original report of prime editing by Anzalone *et al.*^21^, it was found that an 18-bp sequence (“6xHis”) could be inserted at target genomic sites using PE3, a prime editor that achieves high genome-editing activity by nicking both DNA strands. Since the core mechanism of peCHYRON involves the insertion of a 20-bp sequence (*i.e.,* a 3-bp signature mutation and a 17-bp propagator sequence), PE3’s ability to insert 6xHis was a natural starting point in the design of peCHYRON. Several developments were necessary.

First, we reasoned that the modest rates of unintended insertion or deletion mutations (indels) associated with PE3^21^ could be problematic in peCHYRON, because spurious indels at any step of the iterative insertion process would render the recording locus unrecognizable by prime editor and thus terminate recording. We therefore tested whether 6xHis could be inserted using PE2, which does not cut both strands of target DNA^21^ (**Figure 2a**). To our surprise, we found that 6xHis was inserted by PE2 at comparably high efficiency as by PE3. As expected, the 6xHis insertion made by PE2 was accompanied by a much lower rate of unintended insertion or deletion mutations (indels) (**Figure 2a**). Consequently, we decided to use PE2 as the prime editor for peCHYRON.

**Figure 2.**
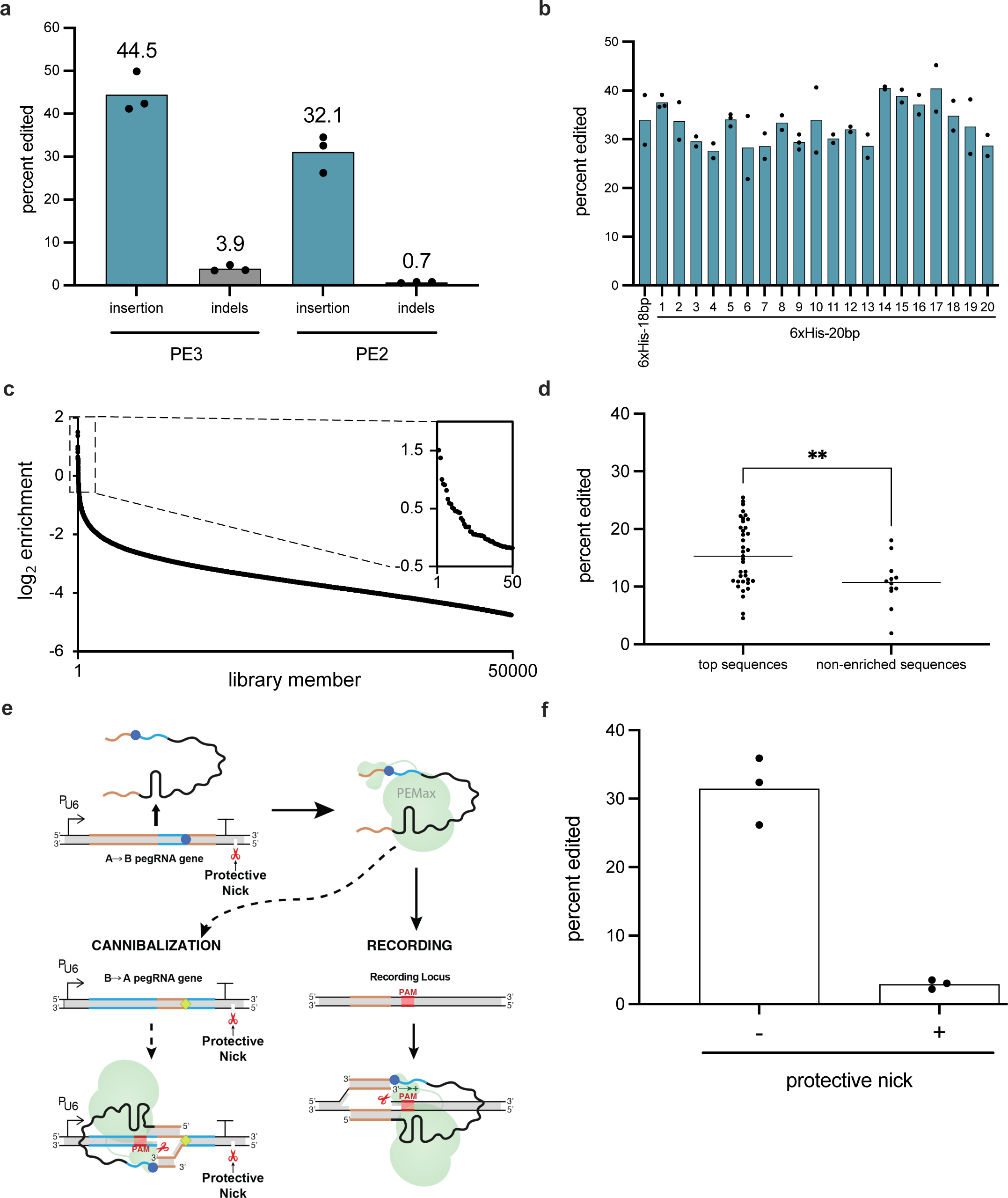
Identifying efficient components for iterative editing with two alternating propagator sequences. **a**, PE2, a prime editing system that does not make double-strand breaks, efficiently adds insertions at the peCHYRON locus. An 18-bp 6xHis sequence was inserted at site3 in 293T cells using PE3 and PE2, and the rates of intended insertions vs. spurious indels were assessed. **b**, 20-bp sequences derived from the 18-bp 6xHis sequence were inserted efficiently. Each 20-bp sequence was nearly identical to the original 6xHis sequence, but with the first 3 bp randomized (3-bp signature + 17-bp propagator sequence). Insertions were made at site3 in 293T cells using PE2. Points represent biological replicates. Efficiencies from different experiments were normalized using samples that were present in each experiment (6xHis-20bp sequences 1, 5, and 9). **c**, Efficient pegRNAs that target 6xHis and insert a different 20-bp sequence (A→B) were rare. A high-throughput screen was used to identify pegRNAs that could alternate with the 20-bp variant of the 6xHis tag to iteratively edit the peCHYRON locus. In the screen, a library of ∼100,000 pegRNAs with variable insertion sequences was transfected into 293T cells along with PE2. High enrichment values correspond to pegRNAs that frequently edited the 6xHis target site without being over-represented in the original transfection mix. The 50,000 library members whose enrichment could be calculated were plotted, and the inset shows the top 50. **d**. Enriched library members directed more efficient editing when tested one-by-one. 37 library members that were among the 0.5% most-enriched pegRNAs and 12 library members that were detected but not enriched were individually cloned and transiently transfected, with prime editor, into 293T cells. Target loci were sequenced after 3 days and editing efficiency quantified. Points show the mean of 3 biological replicates. The difference between the groups was significant by Welch’s t test (P=0.0037). Specific results are shown in Extended Data Figure 1e. **e**, Cartoon showing cannibalization and a strategy for blocking prime editing at the pegRNA genes (*i.e*., cannibalization) while allowing prime editing at the recording locus. The protective nick makes repair favor the unnicked strand and disfavor the nicked, edited strand, thus preventing prime editing. **f**, Prime editing at a genomic locus was blocked by nicking an adjacent site on the same strand. 293T cells were transiently transfected with plasmids encoding PEmax, a pegRNA that directs a 20-bp insertion at the site3 locus, and either an sgRNA that directs a nick at a distant locus (-) or an sgRNA that directs a nick 21 nt 5’ of the cannibalization-associated nick (+), on the same strand. 3 days after transfection, the genomic site was sequenced. The graph indicates the proportion of loci that had been prime edited; each dot represents one biological replicate.

Second, we tested whether the 18-bp 6xHis insertion sequence identified in Anzalone *et al.*^21^ could be converted into a 20-bp insertion sequence in which the first 3 bps constituting the signature mutation are randomizable (**Figure 2b**). A collection of pegRNAs, each of which targets a defined site in the genome (“HEK site3” or “site3”^21,24^) and contains the template for inserting a distinct 3-nt signature mutation followed by a constant 17-nt sequence nearly identical to 6xHis, was tested for PE2-mediated editing in HEK293T (“293T”) cells. Although some variation was observed, most sequences were inserted with high efficiency similar to that of the original 18-nt 6xHis sequence.

Third, we attempted to make a pegRNA that uses the inserted 6xHis-based sequence as its target for subsequent propagation. Recall that the 6xHis-based sequence was inserted at the site3 target. If we consider site3 to be A and 6xHis to be B in peCHYRON’s pair of alternating propagator sequences (see **Figure 1d**), the original pegRNA that installed 6xHis corresponds to the A→B step, and a pegRNA designed to insert site3 at the 6xHis target corresponds to the B→A step. We designed a B→A pegRNA that inserts site3. However, we did not observe efficient editing (**Extended Data Figure 1d**). To find a sequence that could be inserted efficiently, we performed a high-throughput screen in which a library of ∼100,000 pegRNAs, each designed to target the 6xHis site and add a random 20-nt sequence, was transfected into 293T cells that contained the target 6xHis site in their genomes (293T_6xHis_). After 3 days, we sequenced the target sites, obtaining sufficient coverage to assess the efficiency of 50,000 library members, and identified enriched insertions, which we then validated individually for high insertion activity. Efficient insertion sequences were uncommon, but several were identified (**Figure 2c-d, Extended Data Figure 1e**, and **Supplementary Table 1**). For ten of the most efficient insertion sequences, we subsequently optimized PBS length and ensured they could tolerate variable signature mutations (**Extended Data Figure 2a**).

For propagation to occur, an inserted sequence must act as the target for the next round of prime editing. Of the ten efficient insertion sequences, we identified two that tolerated variable signature mutations and behaved as good targets for the insertion of 6xHis sequences containing signature mutations (**Extended Data Figure 2a-b**). Let us call these two sequences from the high-throughput screen B_4_ and B_7_ and the 6xHis sequence A (see **Figure 1d**). The downselection from the screening steps described above ensured that B_4_ and B_7_ could successfully act with A both in the A→B step (*i.e.*, target A and insert B_4_ or B_7_) and the B→A step (*i.e.*, target B_4_ or B_7_ and insert A), providing the necessary ingredients for peCHYRON. When we simultaneously transfected A®B and B®A pegRNA pairs for both B_4_ and B_7_, we observed extended propagation (**Extended Data Figure 2c**). One pair resulted in slightly more efficient propagation (**Extended Data Figure 2c**). Both A®B and B®A components of this pair were able to direct a variety of signature mutations in the original screen (**Extended Data Figure 2d**), so we performed all subsequent experiments with that pair.

## Optimizing peCHYRON for long-term recording

The implementation of peCHYRON requires that all components be delivered to the relevant cells, integrated into their genomes, and maintained over time. Therefore, we created a 293T_6xHis_ cell line into which we integrated all peCHYRON components: a synthetic target site comprised of the 20-bp 6xHis sequence inserted at the genomic site3 locus, prime editor, and A®B and B®A pegRNAs. Ideally, we would see that progressively longer records accumulate at the recording locus, while the genes encoding the recording components remain stable. We observed the accumulation of edits at the recording locus as expected; however, we also observed insertion mutations in the genes encoding the pegRNAs (**Extended Data Figure 3a**). This problem is inherent in the sequence constraints of peCHYRON components – because each pegRNA installs a target site to be used in the next round of prime editing, the sequence of that target site must also be present in the genes that encode the pegRNAs themselves; therefore, A→B pegRNAs can target the B→A pegRNA genes, and vice versa (**Figure 2e** and **Extended Data Figure 3b**). Left unchecked, this “cannibalization” by peCHYRON would destroy the pegRNA genes at the same rate that desired editing occurs at the recording locus. Truly long-term recording would be impossible, as editing efficiency would gradually fall due to depletion of intact pegRNA genes.

To unlock peCHYRON’s long-term functionality, we designed a method to block prime editing at the pegRNA genes without also blocking the desirable editing at the recording locus. Anzalone et al.^21^ showed that, after installing a prime edit, nicking the opposite DNA strand biases host cell DNA repair to maintain the prime edit (PE3). We reasoned that if, after installing a prime edit, we instead nicked the same strand, we would get the opposite result: the host cell would reverse the prime edit. We call this idea “protective nicking.” By targeting protective nicking to a sequence adjacent to pegRNA genes but not present at the recording locus, cannibalization could be selectively blocked without impairing the efficiency of the desired recording (**Figure 2e** and **Extended Data Figure 3b**). To test this idea, we first attempted to block prime editing via protective nicking at an arbitrary genomic site. We transiently transfected 293T cells with plasmids encoding PEmax and a pegRNA directing a 20-bp insertion at the genomic site. To implement protective nicking, we co-transfected a plasmid encoding an sgRNA that would cause nicking near the prime edit, on the same strand (21 nt 5’ of the prime edit). As a negative control, we instead transfected a plasmid encoding an sgRNA that directs a nick at a distant genomic site. When the protective nick was made adjacent to the prime edit, prime editing was reduced ∼10-fold (**Figure 2f**).

We next used protective nicking to prevent cannibalization in a stable peCHYRON cell line over several weeks. To create this stable cell line, we integrated 4 types of PiggyBac cargo plasmids into 293T_6xHis_ cells: one encoding PEmax, marked with mTagBFP2; a second encoding an A→B pegRNA, marked with mCherry; a third encoding a B→A pegRNA, marked with sfGFP; and, optionally, a final plasmid encoding a protective nicking sgRNA. Both A®B and B®A pegRNA plasmids include a target site for the protective nicking guide. An ideal cell line would express all components, including several of each type of pegRNA, and exhibit efficient recording at the intended locus but low cannibalization. To achieve this, for each transfection, a population of cells expressing the peCHYRON components was selected by sorting for high (top 20-30% among positive cells) red, green, and blue fluorescence. We assumed cells that expressed the fluorophore-marked transgenes at high levels were likely to have received high quantities of all transgenes (including the protective nicking sgRNA). These cell lines were passaged for a total of 46 days post-transfection. To ensure that the apparent rate of cannibalization was not artifactually reduced because cannibalization produces longer, harder-to-amplify loci, we amplified a 1kb region of our pegRNA-encoding genes and performed nanopore sequencing to measure cannibalization at multiple timepoints. At 27 days, fewer than 10% of pegRNA-encoding genes were cannibalized in cell lines including the protective nicking sgRNA, but substantial cannibalization had occurred in cells lacking the protective nick (**Extended Data Figure 4a-b**). At later timepoints, the effect of protective nicking was less pronounced, which may be due the fact that we did not select specifically for the protective nicking sgRNA resulting in loss of its expression over time. These experiments showed that protective nicking could block prime editing, not only over a short timecourse (**Figure 2f**), but in the context of a long-term experiment.

Having found a way to maintain the integrity of the pegRNA genes, we used the same experiment to test whether all peCHYRON components could be maintained over time. To test whether the addition of peCHYRON components reduced the fitness of the cells, we set up a competitive growth assay, in which peCHYRON cells (fluorescent in red, green, and blue channels) were mixed with their parent 293T_6xHis_ cells (not fluorescent) in equal cell numbers and passaged as needed. The relative amounts of each cell type were assayed by flow cytometry. We found that the population of cells that were fluorescent in the red, green, OR blue channels (i.e., those that had been engineered) showed very little fitness defect (**Extended Data Figure 4c**). However, the proportion of cells expressing all three fluorescent proteins declined significantly, likely due to silencing of the loci expressing the peCHYRON components, especially PEmax. Indeed, we saw a similar rate of silencing in cell populations that only included peCHYRON cells (**Extended Data Figure 4d**). Rates of silencing and fitness differences of silenced cells likely interact to produce the observed rate of silencing. Although all of our PiggyBac cargo constructs are flanked by insulator sequences, this observation is unsurprising; silencing is a challenge throughout mammalian synthetic biology^25^. To enable long-term stable expression of PEmax, previous works have used constant antibiotic selection^26^ or integration at a safe-harbor locus followed by clonal selection^27^, and some combination of approaches may be required to maintain prime editor expression for recording experiments in animals. We also observed that the cells selectively lost the target site over time (**Extended Data Figure 4e**). Our initial “A” target site was integrated at one copy of site3, which in 293T cells is present in three copies. Therefore, when site3 is subjected to amplicon sequencing it is expected that ∼33% of reads will include the target site. However, this ratio dropped progressively during our timecourse, which is likely explained by prime editing-induced or spontaneous chromosome loss and gene conversion combined with selection against the presence of the target site. Although these observations of component loss over time all posed challenges to peCHYRON, they were still consistent with continued editing (**Extended Data Figure 4f**). Moreover, these challenges are shared with all DNA recording strategies that use prime editor, and likely all strategies that use any CRISPR tool – future work to address them will be paramount.

## peCHYRON accumulates edits over time at genomic loci

Having established that long-term recording is possible, we sought to limit cannibalization and continuously accumulate edits over a longer timecourse. In this experiment, we specifically selected for each component: PEmax (puromycin resistance) and the protective nicking sgRNA (blasticidin resistance) by antibiotic treatment for the first 20 days and pegRNA expression (sfGFP and mCherry expression) by FACS on day 20. This protocol achieved the desired result: almost all recording loci acquired at least one edit by the end of the 66-day experiment, and 75% of pegRNA loci were successfully protected from cannibalization (**Figure 3a**). Indeed, the cells exhibited extensive editing at the recording locus, with edits continuously accumulating for the entire 66-day timecourse (**Figure 3b**). As in the previous experiment, we assessed both cannibalization and recording using long amplicons followed by nanopore sequencing to reduce the effect of each locus’ size on its relative amplification. Consistent with our observation of silencing in **Extended Data Figure 4d**, the rate of edit accumulation, defined as the average increase in edits between time N and time N+1, declined moderately over time (**Extended Data Figure 5a-b** and **Supplementary Table 2a**). At least in two of three replicates, the rates were noisy at later timepoints (**Extended Data Figure 5a**), likely due to some combination of silencing and selection, but the overall picture is one of gradual reduction in rate over the course of the experiment. The rate of edit accumulation declined from ∼4% of loci acquiring an edit per day at day 27 to ∼2% by day 66 (**Extended Data Figure 5b).** Even with this decline, editing rates indicated recording could continue past day 66, and rates are likely to remain more stable when strategies to prevent prime editor silencing are implemented in the future.

**Figure 3.**
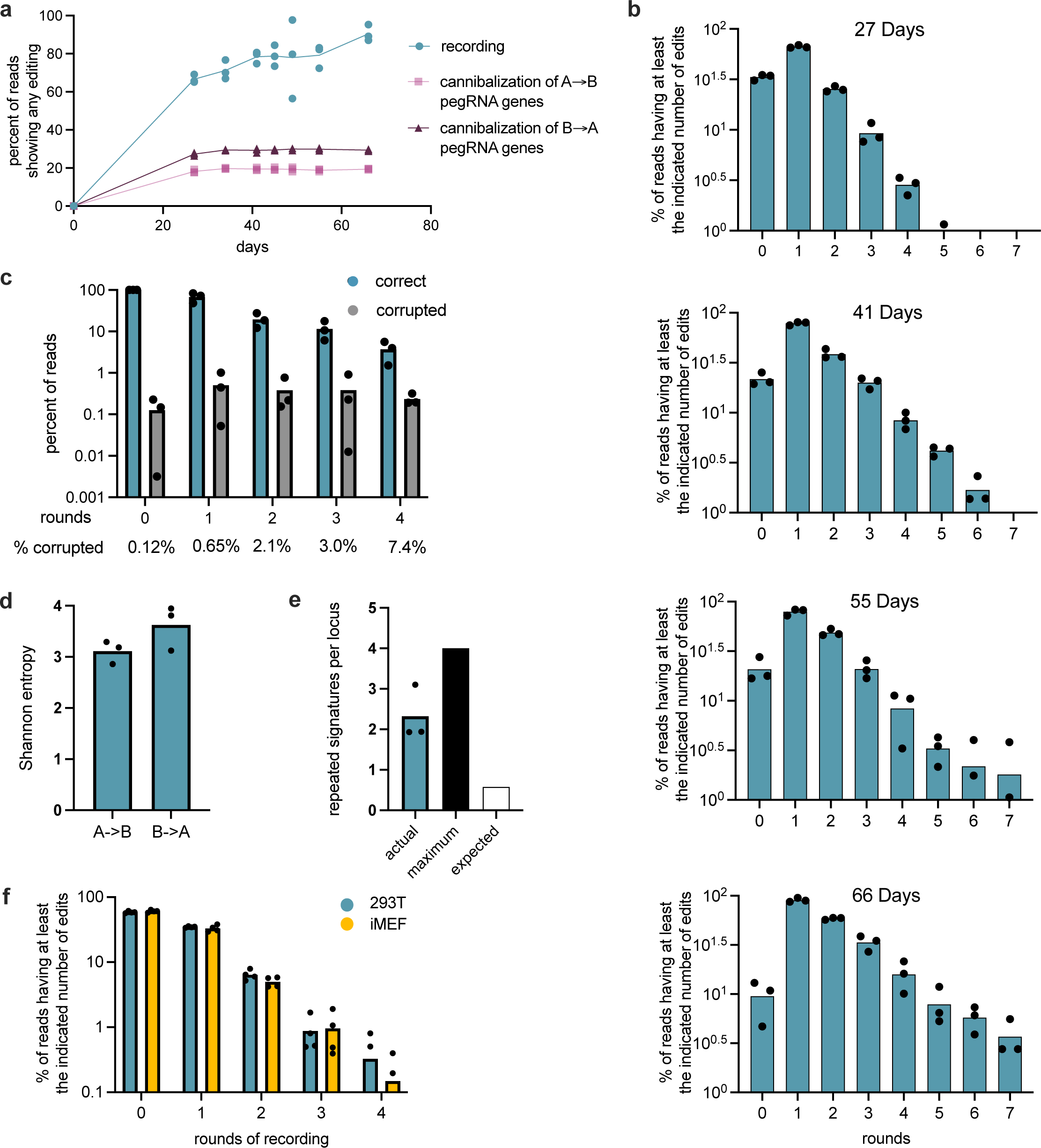
Characterization of peCHYRON performance. **a**, Long-term recording was compatible with limited cannibalization of the pegRNA genes. Expression cassettes for PEmax (marked with puromycin resistance), 21 A→B pegRNAs (marked with mCherry), 21 B→A pegRNAs (marked with sfGFP), and a protective nicking guide, were co-transfected with PiggyBac transposase for stable integration into the genome of 293T_6xHis_. Cells were grown for the indicated time after the initial transfection, then the recording locus, the A→B pegRNA-expressing loci, and the B→A pegRNA-expressing loci were sequenced. For the recording locus, points represent the proportion of reads in which at least one peCHYRON edit occurred. For the pegRNA-expressing loci, any deviation from the original sequence was considered cannibalization and plotted (see Methods). For panels a-e, each point corresponds to one of three biological replicates; some points overlap. **b**, In the experiment shown in (a), at the recording loci, insertions accumulated for at least 66 days. Insertions of 20 bp were considered to correspond to 1 round of insertion, 40 bp to 2 rounds, and so forth. The proportion of loci that had reached at least each number of rounds was plotted, *e.g.*, a locus with an insertion 140-bp long, corresponding to 7 rounds, would be counted towards the total for 1, 2, 3, 4, 5, 6, and 7 rounds. **c**, The rate of spurious indel mutations during 66 days of passaging was low. From Illumina sequencing of the experiment show in a-d, each read was assigned to a bin according to its size; insertions that exactly match the expected sizes were considered “correct,” and all other mutations were considered “indels.” The correct totals plotted include all reads that are that size or above, as in b. The % corrupted value was determined by dividing the corrupted value by the correct value. We note that the proportions of correct loci shown differ between this panel and the bottom panel of b, even though these panels show data gathered from the same samples. This difference is explained by the library prep strategies. In peCHYRON, each round of recording increases the length of the locus by 20 bp. In our nanopore sequencing pipeline, used for all assessments of recording efficiency, the recording loci were sequenced as part of a 2.3-kb amplicon, so 20-bp differences are not likely to significantly affect the amplification rate. The error rate of nanopore sequencing is too high for accurate assessments of corruption, so we needed to use Illumina sequencing. For Illumina sequencing, we used an amplicon that is 154 bp for loci with no insertions. Thus, 5 rounds of recording increased the size of the PCR product by 65%. It is not surprising, therefore, that we observed an underrepresentation of longer loci (**Extended Data Figure 5d**). In addition, due to read length restrictions, we were only able to sequence 108 bp of inserted sequence, so corruption could only be assessed for the first four rounds. These considerations should not affect the ratios of correctly- and incorrectly-sized loci, which reflected very low rates of corruption: 0.65% of loci at round 1 gradually increased to 7.4% by round 4 **d**, Within each population of cells, diverse signatures were incorporated. Shannon entropy was calculated (see Methods) for A→B and B→A edits after 66 days of recording. **e**, Within individual cell lineages, signature diversity was more limited. For all loci that had undergone 6 rounds of recording, the number of repeated sequences within a single locus was calculated, and the average plotted, along with the maximum number of repeats, and the expected number, given the Shannon entropy values calculated in c (see Methods). **f**, peCHYRON recording was similarly efficient in multiple cell types. peCHYRON recording cell lines were created from 293T and SV40-immortalized mouse embryonic fibroblasts in parallel. Recording loci were sequenced 15 days after transfection. Two technical replicates for each of two biological replicates are shown.

Aside from maintaining editing rates over time, long-term recording also requires that editing remains efficient as the recording locus accumulates edits (*i.e.*, the target site must not lose its receptiveness to editing as a result of past edits). To test whether this is the case for peCHYRON, we used the data from the 27- and 34-day timepoints and calculated the frequency of observing an additional edit during the intervening week as a function of the number of edits already present at day 27. To simplify analysis, we assumed that each locus would receive no more than one edit during that week (see **Supplementary Table 2a** for detailed calculations). The rate of additional editing trended higher for sequences that had more previous edits (**Extended Data Figure 5c**). We cannot exclude the possibility that cells with more edits at day 27 had higher prime editor expression, which would then promote further editing and contribute to our observation of higher editing frequencies for longer records. Nonetheless, the lack of any major efficiency declines as a function of previous editing at the recording locus, in contrast to precipitous efficiency drops observed after few successive edits at the same recording locus in recorders that rely on DNA double-strand breaks^10,11,13,14,18^, should promote peCHYRON’s continuous operation.

We next considered another parameter relevant to long-term recording: “corruption,” or how often a round of recording led to an unintended edit at the recording locus. Because unintended edits often terminate recording and erase previous edits, and corruption would progressively accumulate over rounds of recording, even a moderate rate of corruption per round (*e.g.*, 10%) would be incompatible with long-term recording. We resequenced the final timepoint from our experiment above using an Illumina pipeline. Corruption was assessed as the rate of insertions of unexpected size. We found the amount of corruption was low, with only 0.65% to 7.4% of sequences receiving 1 to 4 rounds of sequential editing becoming corrupted (**Figure 3c**). This means that after enough time has passed for many loci to experience 4 rounds of editing, 92.6% of loci remain uncorrupted and available for continued faithful recording.

In addition to continuously creating and maintaining records, peCHYRON must install different signature mutations with relatively little bias in order to record unique cell lineages or many inputs in parallel. The timecourse above, in which 21 pegRNAs for each type of edit were included, allowed us to calculate the information content of each edit across each whole polyclonal population. The observed Shannon entropy was 3.1 bits per A®B edit and 3.6 bits per B®A edit (**Figure 3d** and **Supplementary Table 2b**), which showed an acceptably low bias. For example, if 21 possible edits were incorporated with no bias, we would expect a Shannon entropy value of ∼4.4 bits^28^. When we examined the sequences of all loci in the experiment that had encoded 6 signatures (the longest we could sequence with our Illumina pipeline), we observed more repeated signatures than would be expected based on the Shannon entropy values calculated for the entire population (**Figure 3e** and **Supplementary Table 2c**). This observation suggests that each individual cell expresses only a few pegRNAs highly. The number of pegRNAs expressed per cell could be increased by delivery of pegRNAs with viruses at high multiplicity of infection, as used for DNA Typewriter^15^, or by encoding multiple pegRNAs per cargo plasmid. Thus, peCHYRON encoded information supplied by pegRNAs with limited bias; more information will be encoded in each cell with improvements in the making of cell lines.

## peCHYRON accumulates edits in multiple cell types

All experiments described so far were performed in 293T cells, but the application of peCHYRON will likely be in cancer and developmental biology, requiring that the system operate in many different types of cells. We tested the efficiency of peCHYRON recording in an additional cell type. In parallel, peCHYRON cell lines were constructed by PiggyBac transposition in 293T cells and mouse embryonic fibroblasts immortalized with SV40 large T antigen (iMEFs) (**Figure 3f**). Cell lines were selected by treatment with puromycin (on PEmax plasmid) and blasticidin (on nicking guide plasmid), but no other sorting was performed. At 15 days post-transfection, both cell lines showed comparable editing efficiency (**Figure 3f**). This similarity suggests that our characterization of peCHYRON recording characteristics will apply to cell types in which peCHYRON could be used to discover new biology.

## Reconstruction of lineage relationships using peCHYRON

We applied peCHYRON to trace the lineage relationships among populations of cells descended from a parent population *via* a complex splitting process (**Figure 4a**). To achieve the maximum possible editing efficiency, we used serial transient transfections to drive high levels of prime editor and pegRNA expression. We transiently transfected one population of 293T_6xHis_ cells with plasmids encoding PE2 and 42 pegRNAs representing A®B and B®A pairs with varied signature mutations. After allowing the cells to grow for 8 days, we isolated 4 populations of 20,000 cells each. The next day we transfected again, allowed the cells to grow for 3 days, then split each population to yield 8 populations. Cells were transfected again the next day, then allowed to grow for 4 additional days, before being split again to yield 16 final populations. Two days later, ∼1.6 million cells were collected from each population, DNA was extracted, and the peCHYRON recording locus was subjected to amplicon sequencing. Three of the 16 populations sequenced poorly, which we took as an opportunity to exclude from analysis so that the researchers performing the reconstruction were blinded to both the identity of the wells and the exact shape of the lineage tree. Sequences in the remaining 13 populations were first filtered by length (number of insertions), then by frequency of occurrence. We discarded all sequences with fewer than 4 rounds of insertion as the probability of generating a specific sequence of 3 signature mutations by chance rather than by descent was significant. We discarded sequences with frequencies of occurrence below a cutoff determined by the kneedle algorithm^29^ to remove spurious sequences that were likely library prep artifacts. After these two filtering steps, a total of 4571 sequences remained (**Supplementary Table 2d**). For each pair of wells, we counted the number of shared identical peCHYRON sequences to calculate Jaccard similarity^30^ and then used the Jaccard similarity scores to perform agglomerative hierarchical clustering. The resulting tree accurately reconstructed all aspects of the splitting procedure (**Figure 4b**).

**Figure 4.**
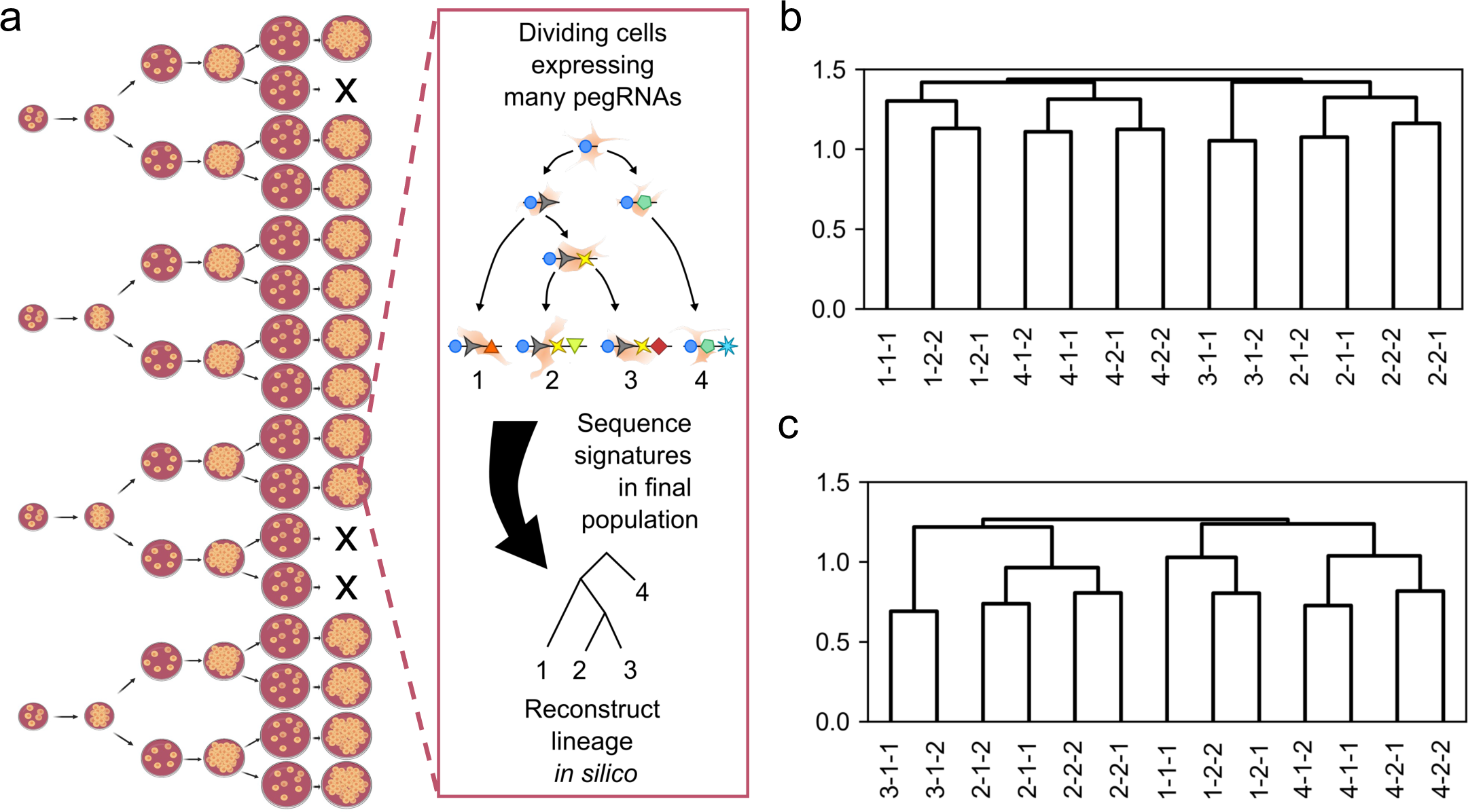
peCHYRON enables cell lineage tracing. **a**, Culture splitting procedure for a lineage tracing experiment using 293T cells transiently transfected with peCHYRON components. Inset shows how signature mutations accumulate in individual cells, then inform lineage reconstruction. **b**, Culture splitting patterns were accurately reconstructed using the Jaccard similarity index. **c**, Culture splitting patterns were accurately reconstructed using a modified Jaccard similarity index that accounts for sequences that share early signature mutations but diverge in later signature mutations.

Calculating Jaccard similarity worked well for comparing populations of cells, but it only took into account identical peCHYRON sequences shared among the populations. Because the peCHYRON recording locus progressively acquires signature mutations in temporal order, related cells can share initial edits and then diverge. Sequences with shared early signatures but diverged late signatures can provide higher resolution information about lineage relationships, theoretically approaching single-cell-resolution. We sought to establish a lineage reconstruction algorithm that would take advantage of this ordered nature of recording. To do so, we modified the Jaccard similarity index to allow for partial matches. Pairs of populations were compared as follows. First, each sequence of signature mutations was split into a collection of all possible subsequences that started from the first signature mutation (*i.e.,* prefixes). Each prefix generated from a sequence was given a fractional weight equal to the inverse of the number of prefixes making up the sequence. This resulted in a multiset of prefixes. We then computed a weighted multiset Jaccard similarity between all pairs of multisets. To illustrate this, consider a record ABCD where A, B, C, and D each represent a unique 3-bp signature mutation. This is split into 4 prefixes, A, AB, ABC, and ABCD, each given a weight of ¼. Now, consider a second record, ABCEF. This is split into 5 prefixes, A, AB, ABC, ABCE, and ABCEF, each given a weight of ⅕. When these are compared, the prefixes A, AB, and ABC match, and we add the minimum count, ⅕, for each match. The sum of these matches becomes the numerator for the Jaccard similarity, and the denominator is simply the sum of the cardinalities of the sets minus the numerator. With this new algorithm, the full splitting procedure was again accurately reconstructed (**Figure 4c**). Lineage reconstruction with this prefix Jaccard algorithm outperformed that done with traditional Jaccard similarity at almost every level of random downsampling and gave near-perfect reconstructions even when downsampling to only 20% of the data (**Extended Data Figure 6**), forecasting the utility of this reconstruction heuristic in realistic lineage tracing experiments where populations are poorly sampled.

## Reconstruction of cellular event histories using peCHYRON

One of the major promises of ordered recording is tracing sequential patterns of signaling that underlie the acquisition of specific cell fates. Ultimately, we would wish to read out the signaling history of single cells. A near-future goal would be to perform simultaneous event recording and single-cell transcriptomics, identify sub-populations based on their transcriptomes, then use the recorded sequences of all the cells in a sub-population to understand the history of that sub-population. A more immediately achievable goal is to treat separate populations of cells as sensors of their environment over time by creating DNA recorders that make specific records in response to specific environmental signals. Ordered recording at CRISPR arrays in both bacterial populations^5,8,31^ and single bacteria^32^ has been used to successfully report orders of signals experienced by the cells. To test a similar capability in mammalian cells, we used peCHYRON to record, then reconstruct, the order of transient events experienced by populations of 293T cells. Specifically, two types of transient events were recorded. In one set of experiments, we aimed to record the order of various pegRNAs delivered to cells. In the second set of experiments, we aimed to record the order of chemical signals to which cells were exposed. In both cases, histories could be reconstructed with high fidelity by reading out peCHYRON sequences (**Figure 5** and **Extended Data Figures 7-10**).

**Figure 5.**
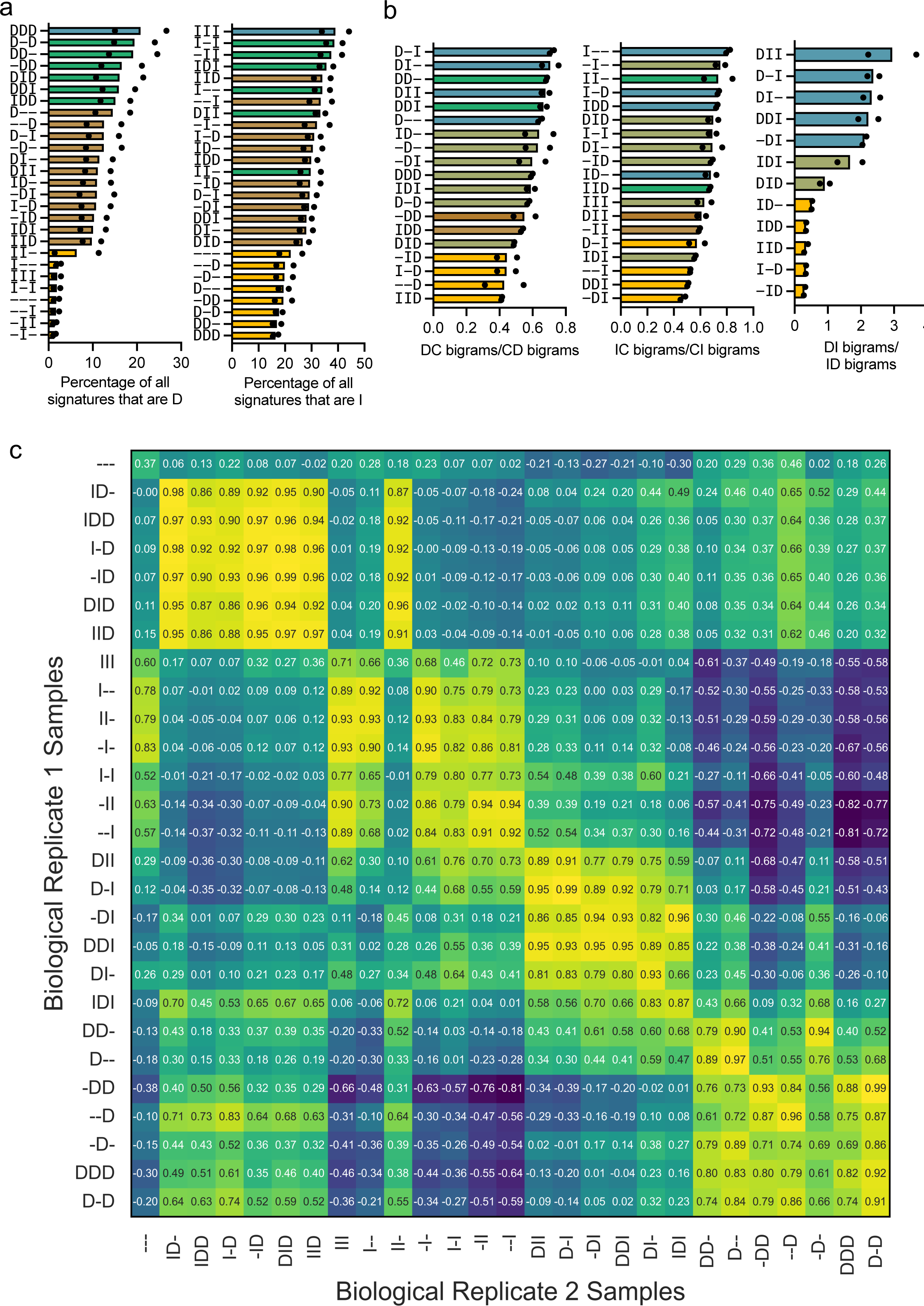
peCHYRON enables temporally-resolved event recording. **a**, 293T_6xHis_ cells in which PEmax, the protective nicking sgRNA, TetR, LacI, and pairs (A→B and B→A) of constitutive or inducible pegRNAs were stably integrated, then exposed to doxycycline (dox, indicated by the D signature), IPTG (I signature), or no inducer (constitutive, or C, signature only) for 3 epochs of 12 days each. After the 3^rd^ epoch, recording loci were sequenced and the relative abundance of D unigrams (left panel) or I unigrams (right panel) was calculated. Dots indicate the value for each of two biological replicates. Bars are arranged in rank order and colored according to the actual duration of chemical treatment, *i.e.*, three, two, one, or zero epochs. **b**, Ordered pairs of recorded signatures (extended bigrams, see Extended Data Figure 7b) can be used to accurately determine the overall order of chemical exposures. For samples exposed to dox, the ratio of DC to CD bigrams was calculated (top panel). An analogous procedure was done for IC to CI bigrams (middle panel) and DI to ID bigrams (bottom panel). Dots indicate the value for each of two biological replicates. Bars are arranged in rank order and colored according to the actual timing of the chemical treatment (top and middle) or the relative timing of dox and IPTG treatment (bottom panel). In the top and middle panels, bars are colored differently if treatment occurred in the first epoch only, first and second epoch only, second and third epoch only, third epoch only, or distributed evenly throughout the timecourse (*i.e.*, all epochs, first and third epochs, or second epoch only). In the bottom panel, bars are colored differently if dox preceded IPTG, IPTG preceded dox, or both. **c**, Unknown histories of cell populations can be inferred by comparing peCHYRON loci sequences to model populations with known histories. For the experiment also shown in a-b, the signature mutations were extracted from peCHYRON records, then the sequence of signature mutations from each recording locus were computationally stretched to a constant length (this is 60 because it is the least common multiple of all insertion lengths in the data set). Stretched records were pooled together in a manner that allows each insertion length to contribute equally to the final pool long insertions are weighted more heavily to prevent being overshadowed by short, abundant insertions that are less informationally rich (*i.e.*, records of each length were weighted by the inverse of their abundance in the total pool). At every position along the stretched records, the proportions of each signature were calculated. The fold change of this proportion relative to a control in which cells were exposed to no inducer for 18 days was plotted for dox (D, green) and IPTG (I, red). These stretched plots are shown in Extended Data Figure 10. All samples in biological replicate 1 were compared to all samples in biological replicate 2 by computing the Spearman coefficient between the stretched record for each of D and I. For each pair, the Spearman coefficients for D and I were averaged to construct a Spearman Coefficient matrix. Hierarchal (agglomerative) clustering was performed on the coefficient matrix and represented as a heatmap.

To interpret peCHYRON records, a previously reported bigram method^15^ and a novel “signature stretching” method were employed (**Extended Data Figure 7a-b**). To illustrate these approaches, consider the simple case of two signals, X and Y, each of which promotes expression of a set of pegRNAs that install a signature X or Y. The order of signature mutations recorded in peCHYRON loci can reveal the order in which X and Y occurred, since each signature mutation reflects the identity of an exact pegRNA, and the sequential nature of insertion accumulation preserves temporal information. In many cases, the bigram method is appropriate for interpreting the order of signals. For this, the frequency of ordered pairs of signatures is calculated across all recording loci in a population of cells. The frequency with which X appears before Y, or vice versa, in individual records, should correspond to the order of the signals: if X occurred before Y, we would expect the bigram XY to be more abundant than YX (**Extended Data Figure 7b**). Although bigram frequencies have been previously reported to successfully recapitulate the order of pegRNA pulses^15^, an inherent limitation of this approach is that it cannot resolve repeated pulses of a given signature. For example, if a population of cells experienced signal X, then Y, then X again at a later time, the bigram frequencies would show high incidence of both XY and YX bigrams such that signal progressions following the order X-Y-X would be difficult to distinguish from those following the order X-Y-X-Y-X, etc. To overcome this problem, we developed another analysis approach that leverages the relationship between timing of signals and position in the peCHYRON recording locus. All records with 2 or more insertions contain some information about the true order of signals experienced, so we aggregate this information by computationally “stretching” the signatures from all such records to the same length, then calculate the fraction of each signature at each position among the pool of stretched records. We then calculate the enrichment of each signature relative to controls with no signals present. For example, if all records are stretched to a length of 60, a record with two insertions with signatures XY would be stretched to 30 Xs followed by 30 Ys. A record with three insertions XYX would be stretched to 20 Xs, then 20 Ys, then 20 Xs (**Extended Data Figure 7b**). Since longer insertions are intrinsically more informative because they originate from cells that recorded signals more frequently throughout the course of the experiment, records are weighted proportionally to their length.

We first validated peCHYRON’s ability to perform temporally resolved event recording by reconstructing pulsed expression of pegRNAs delivered to a population of cells. For this, we used 293T_6xHis_ cells into which PEmax and the protective nicking guide had been stably integrated. We split this cell line into 2 separate populations, then used a random number generator to choose a pair of A®B and B®A pegRNAs to transfect into each population. Both members in the pair of pegRNAs install the same signature mutation as each other. We repeatedly chose random pegRNA pairs to transfect, for a total of 4 rounds of sequential transfections. Once used, a pegRNA pair (and their signature) was never repeated. We collected cells at the end of the experiment, extracted DNA, and subjected the peCHYRON locus to amplicon sequencing. As expected, because each signature mutation was only used once, the order of transfections in our experiment was correctly inferred by calculating the frequencies of all extended bigrams (**Extended Data Figure 7c**). For the first transfection, the corresponding signature was only found in the first position of bigrams, while all signatures corresponding to all three following transfections appeared in the second position. Signatures introduced in the second transfection were only followed by signatures introduced in the third and fourth transfections, and so forth. The signature introduced in the last transfection was never found in the first position in an extended bigram. When the records were computationally stretched so that proportions of each signature at each position could be calculated, the true order of transfections was again reconstructed (**Extended Data Figure 7d**), as the first appearance of each signature on the stretched graph reflected the order of transfections.

We also created cells that could sense and record patterns of chemical signals, without a need for interventions such as transfection. We made cell lines that express PEmax and a protective nicking guide, as well as three different pairs of A®B and B®A pegRNAs with different, known signatures. One pair expressed constitutively, one pair expressed in response to doxycycline (hereafter dox), and one pair expressed in response to IPTG. All necessary constructs were transfected as PiggyBac cargo along with a transient PiggyBac transposase plasmid. Integrants were selected with puromycin, which marks PEmax, and blasticidin, which marks the protective nicking sgRNA. Six days after the transfection, cells were passaged into medium containing dox, IPTG, or no drug. We completed a total of 3 epochs of 12 days each. We tested 27 exposure patterns, representing all possible combinations of dox exposure, IPTG exposure, and no treatment, with two biological replicates of each exposure pattern. Records were read out by amplicon sequencing of DNA from 50,000-150,000 cells per sample.

We first asked whether the data from the 27 samples in each replicate could be used to determine the total amount (*i.e.,* zero, one, two, or three epochs) of dox and IPTG to which each sample had been exposed. We calculated the proportion of signatures that corresponded to dox (“D” signatures) or IPTG (“I” signatures) exposure in each sample. In all cases, a rank ordering of samples by D signature proportion ordered the samples by actual dox exposure duration (**Figure 5a**). Rank ordering by I signature proportion ordered 24/27 samples by actual IPTG exposure duration (**Figure 5b**), consistent with the IPTG-inducible promoter’s much higher leakiness (**Figure 5a-b**). We also tested whether the different durations of chemical exposure could be determined from the data with no prior knowledge of how many groups there were. For all the samples within each replicate, we performed k-means clustering on either D (**Extended Data Figure 8a-b**) or I (**Extended Data Figure 8c-d**) proportions using a script that automatically determines the number of clusters by optimizing for both low inertia and fewer clusters. In all cases, the data clustered into 3 groups, implying three durations of exposure to D or I inducers. In reality, there were four durations of exposure for either D or I inducers – no inducer, one epoch of inducer, two epochs of inducers, and three epochs of inducer. However, there is only one sample that would correspond to three epochs of inducer in contrast to several samples for each of the other three categories, so it is understandable that the one sample group would cluster with another group. Also in each case, the sample treated with the inducer for the first epoch was erroneously clustered with the samples exposed for two epochs, likely reflecting the higher activity of the recorder at the beginning of the experiment, before any silencing could occur. There were a few errors in the I clustering in Replicate 2 (**Extended Data Figure 8d**), and the sample “II-” in Replicate 2 was clustered with samples exposed to dox for one epoch (**Extended Data Figure 8b**), likely reflecting experimental error. Otherwise, the duration of samples’ chemical exposure was correctly reconstructed.

We next asked whether the relative timings of signals and no treatment could be resolved similarly and reasoned that ratios of bigrams including I or D and the constitutive signature “C” could reveal whether I or D was added early or late. These ratios correlated weakly with the actual timing of dox and IPTG treatment (**Figure 5b** and **Extended Data Figure 9a-d**). In contrast, the relative ordering of dox and IPTG exposure was reliably reconstructed by computing the relative proportions of DI and ID bigrams. The rank ordering of samples by this proportion exactly corresponded to the true order of dox and IPTG treatment, *i.e.,* samples in which dox treatment preceded IPTG treatment had a high DI/ID ratio (above 1), samples in which dox and IPTG exposures alternated had a ratio around 1, and samples in which IPTG preceded dox had a low DI/ID ratio (below 1) (**Figure 5c**). In an ideal case, these three groups would be put into three clusters. When k-means clustering was performed, 11 of 12 determinations were correct for Replicate 1 (**Extended Data Figure 9e**) and 12 of 12 for Replicate 2 (**Extended Data Figure 9f**). Thus, simple analyses of peCHYRON loci can reconstruct several aspects of the history of populations of cells, especially the dose of a signal and relative timings of chemical exposures.

Because exposure patterns that included repeated instances of each signal (or lack thereof) were more difficult to resolve, we hypothesized that they could benefit from our analysis method that computationally stretches the records. When the records were computationally stretched, the proportion of edits at each position, compared to a no signal control, again successfully reflected the order of the dox and IPTG treatments for all samples (**Extended Data Figure 10**). In addition to the relative order of signal exposures, the absolute timing of signal was also captured in the records. For example, samples that were exposed to dox in the first or second epoch showed enrichment for the corresponding signature in the early or middle positions of the stretched records, whereas samples that were treated with doxycycline in the third epoch exhibited a later increase in the associated signature. The same was true for IPTG treatment. In addition, the relative durations of induction applied to each sample were often distinguishable, allowing for comparisons of cells’ experiences among samples in the same experiment. Although the editing outcomes for each pattern of signal progressions followed expected trends, we acknowledge that the stretched records for certain similar epoch patterns were indistinguishable in some cases (*e.g.*, two epochs of IPTG and three epochs of IPTG). In the future, higher-efficiency editing by peCHYRON will enhance its ability to distinguish similar but not identical patterns of cell history.

For use of peCHYRON to record transient cellular events in an animal model, a method for interpreting records that would be able to leverage all of the data contained in sequences at the recording locus would be to perform experiments in cell culture with known durations and known orders of the events, then compare those cell culture records to the records obtained from animal experiment. To make it possible to determine to what extent the current version of peCHYRON could be used in this way, we compared, for all 27 possible signal progressions, the stretched records from one experiment (Replicate 1) to a repeat of that experiment (Replicate 2) where the Replicate 2 experiment used cells containing 10% less PEmax plasmid added when creating stable cell lines, mimicking a slightly different cell type or state. We computed the Spearman correlation between the weighted compositions of stretched records from each signal progression from Replicate 1 and that from Replicate 2. We reasoned that if a given signal progression’s Replicate 1 record most closely correlated with the same signal progression’s Replicate 2 record and vice-versa, this would mean that (1) records reproducibly reflect the signal progression responsible for their generation and (2) that the signal progressions are distinguishable from each other, here among the set of all 27 possible progressions following the three-epoch schedule. We found that some but not other aspects of the possible progressions were reproducibly distinguishable from each other (**Figure 5c**). For 44% of the samples in Replicate 1, the highest correlation was to the corresponding sample in Replicate 2. For an additional 33%, the highest correlation was to a sample that shared two of three epochs. Hierarchical clustering revealed the epoch patterns that drove similarities between stretched records. Five clusters formed: samples with no treatment, samples in which IPTG preceded dox, samples in which IPTG but not dox was added, samples in which dox preceded IPTG, and samples in which dox but not IPTG was added. Thus, stretched peCHYRON records reliably report which of two types of chemical exposures occurred, and their timings relative to each other, as well as some information about the absolute timing and dose of chemical exposures.

Taking into account all analyses, an experiment in which 27 patterns of chemical exposures, including no treatment, must be distinguished from each other by sequencing peCHYRON loci is at the edge of peCHYRON’s current capabilities. peCHYRON performed very well when asked to distinguish the duration of chemical exposure, limited only by reporter quality: reconstruction of dox treatment duration was near perfect (**Figure 5a** and **Extended Data Figure 8a-b**), whereas a leaky reporter caused some but not many errors in reconstruction of IPTG treatment duration (**Figure 5a** and **Extended Data Figure 8c-d**). The relative timing of dox and IPTG treatment can be reconstructed nearly perfectly (**Figure 5b-c** and **Extended Data Figure 9e-f**), but the absolute timing of chemical treatment was more challenging to reconstruct. Patterns in which dox or IPTG was added only in the first or only in the last epoch were always distinguished from each other (**Figure 5b** and **Extended Data Figure 9a-d**), but more intermediate patterns were often confused. These shortcomings will be addressed in the future by engineering better reporters and more efficient recording.

## Discussion

Our results establish peCHYRON as an advanced DNA recorder that autonomously generates ordered, high-information records inside cells. These records are exceptionally durable and can be parsed to decode the temporal order of recorded events. They are also open-ended; that is, records can be arbitrarily long since each round of recording installs a target site for the next round. In keeping with these favorable performance characteristics, peCHYRON was used to accurately reconstruct complex cell lineage relationships and event histories in proof-of-concept experiments that exploit the information available in temporally ordered records.

For a DNA recorder to continuously operate in the context of a developing organism, it needs to perform robustly in a variety of cell types, as the cells that are being tracked undergo phenotypic changes. One major source of variation in genome editing outcomes between cell types arises from each host cell’s complement of DNA repair proteins^33^. Prime editing is enabled by widely-expressed DNA replication factors^23,34,35^ and can be antagonized by mismatch repair (MMR)^23^ and double-strand break (DSB) repair. DSB repair (*e.g.*, non-homologous end joining and homologous recombination) can lead to unintended indels when both DNA strands are nicked^36^, as in some prime editing strategies (*i.e.,* PE3^21^). However, because only one strand is nicked during prime editing with the PE2 architecture used by peCHYRON, we do not expect the DSB repair pathway to be triggered during recording. In agreement with this model, very low rates of unintended indels are created with PE2 (^21^, **Figure 2a**, and **Figure 3c**). The other known pathway that typically curbs prime editing, mismatch repair (MMR), is deficient in 293T cells^37^. Thus, prime edits that are highly efficient in 293Ts often perform substantially worse in other cell types^23^. To peCHYRON’s benefit, however, insertions longer than 13 bp are not subject to MMR, even in MMR-competent cells^38^, meaning our 20-bp insertions are unlikely to be affected. The efficiency of smaller prime edits that are subject to MMR has been increased by expressing a dominant negative mutant of *MLH1*^23^, but implementing this MLH1dn strategy in an animal model would likely interfere with the biology we seek to study. Due to its minimal interface with host DNA repair, peCHYRON is exceptionally well-positioned for consistent activity across cell types, without disrupting the native DNA repair milieu in the cells of interest.

It is instructive to compare peCHYRON to DNA Typewriter^15^, which was pre-printed simultaneously with peCHYRON and has subsequently been published (**Supplementary Table 3**). DNA Typewriter uses prime editor to install 6-bp insertions in temporal order on pre-defined recording arrays. Key advantages of DNA Typewriter over peCHYRON include relative simplicity of optimization and part expression (*i.e.*, only one type of pegRNA and no protective nicking guide), and high efficiency (*e.g.*, in a lineage tracing experiment in a 293T clonal cell line, the majority of loci recorded at least four edits)^15^. Key advantages of peCHYRON over DNA Typewriter include evasion of MMR, and the ability to add its own target sites^9^, allowing in principle arbitrarily many rounds of recording and opening up the possibility of editing at many sites in parallel in highly repetitive mammalian genomes^9^ or evading target site loss (**Extended Data Figure 4e**) by optimizing recording location. Stable recording for many rounds per site, at many sites per cell, will be necessary to track the activity of many pathways in parallel at single-cell resolution. In addition, peCHYRON is likely to have an advantage in contending with the inherent instability of highly repetitive DNA arrays^15,39^, since repetitive units are added only as records made by peCHYRON rather than being present throughout the experiment as in DNA Typewriter. Further, the periodicity of repeating sequences in peCHYRON is 40 bp, compared to 14-20 bp for DNA Typewriter, so that acquisition of an equal number of edits with peCHYRON and DNA Typewriter would result in ≥ 2-fold more repeated sequence motifs in DNA Typewriter than in peCHYRON. The peCHYRON architecture represents a useful addition to the DNA recording toolkit, especially for applications that call for indefinite expansion of the recording locus (*e.g.*, recording over very long timespans).

In the near term, we aim to improve the resolution of recorded lineage phylogenies and the confidence of event history reconstructions by increasing the efficiency of editing with peCHYRON. We may achieve this *via* optimizing the propagator sequences^38^ and taking advantage of the ever-growing prime editing toolkit (e.g.,^27,40^). However, even with the current editing efficiencies, the central architecture of peCHYRON possesses the critical features of a DNA recorder well-suited to long-term experiments. For example, deep lineage tracing on the scale of a whole mammal requires a recorder that can continuously operate throughout development and generate durable records capable of distinguishing among billions of cells^41^. peCHYRON can already continuously operate for at least 2 months with relatively small decreases in the rates of propagation, and, at least across a population of cells, it encodes 3.1-3.6 bits of information in each sequentially inserted record (**Supplementary Table 2b**) such that a record containing just 11 signature mutations has ∼40 billion possible states, similar to the number of cells in an adult mouse. peCHYRON is exceptionally well-matched for temporally-resolved multiplexed signal recording, as it accumulates records in an ordered manner, and the number of signals it can record can be expanded by expressing pegRNAs with known signature mutations in response to promoters of interest. Here, we have recorded the orders of chemical signals added to cell populations and have demonstrated our pegRNAs can tolerate at least 41 unique 3-nt signature mutation sequences (**Supplementary Table 2b**), each of which could theoretically be linked to an additional inducible promoter to record highly complex histories of what cells experience *in situ*. Applications of peCHYRON beyond the proof-of-concept experiments shown here will capitalize on these and other opportunities in the quest to understand the detailed histories of individual cells in animal biology and development

## Acknowledgements

We thank Christine Duong and Seanjeet K. Paul for technical assistance; Luke Koblan, Jingyi Jessica Li, Kamen Simeonov, members of the Liu Laboratory, Olga Razorenova, and Jordan Woytash for helpful discussions; and David Liu (Broad Institute), Timothy Lu (MIT), and Hana El-Samad (UCSF) for plasmids. This work was funded by NIH grants R35GM136297, DP2GM119163, and R21GM126287 to CCL; NIH grants K99GM140254 and R00GM140254, and start-up funds from Rice University to TBL; NSF GRFP and AHA Predoctoral Fellowships to CKC; and a fellowship from the NSF-Simons Center for Multiscale Cell Fate Research (NSF Award 1763272) to TBL. VJH is supported by Medical Scientist Training Program grant T32-GM008620. Sequencing on the Illumina MiniSeq was performed by the Genetic Design and Engineering Center at Rice University, which receives financial support from the Cancer Prevention and Research Institute of Texas (Award # RP210116).

## Author contributions

TBL, CKC, and CCL conceived the project. TBL, CKC, and CCL designed experiments. TBL, CADH, SKR, CKC, GL, MF, AS, and MJPC performed experiments following protocols developed by TBL, CKC, and CADH. TBL, CKC, VH, MCD, AM, and CCL designed and performed analyses. VH, CKC, TBL, MCD, and MJPC wrote code. TBL, CKC, and CCL wrote the paper, with input from all authors. CCL and TBL procured funding and oversaw the project.

## Supplementary Material Contents

Supplementary Tables

Supplementary Table 1. Detailed results from screen shown in **Figure 2c**.

Supplementary Table 2. Data and entropy calculations underlying **Figures 3-4**.

Supplementary Table 3. Comparison between peCHYRON and DNA TypeWriter.

Supplementary Table 4. Guide to plasmids used, HTS datasets available at the NCBI Sequence Read Archive, HTS primers, and cell lines.

## Methods

### Plasmid cloning

Polymerase chain reactions (PCRs) were performed with Q5 Hot Start High-Fidelity DNA Polymerase (NEB M0493) or Phusion Hot Start Flex DNA Polymerase (NEB M0535). PCR reagents and all DNA-modifying enzymes were also purchased from NEB, and all primers were synthesized by Integrated DNA Technologies (IDT). Plasmids were made by standard Gibson assembly^42^, *in vivo* recombination, Golden Gate assembly, or NEBuilder HiFi assembly (NEB E2621). Specifically, pegRNA-expressing plasmids were made either by adding or changing sequences on PCR primers, then performing *in vivo* recombination in Top10 bacteria, or by ordering pegRNA sequences as long oligos from IDT, then performing NEBuilder HiFi assembly. PiggyBac cargo plasmids were assembled by Golden Gate. All plasmids to be used for transfection were purified from Top10 or NEBStable *E. coli* (NEB C3040) with HP GenElute Midi or Mini kits (Sigma # NA0200 and NA0150).

Plasmids encoding PE2 (Addgene plasmid #132775^21^), PEmax (Addgene plasmid #174820^23^), and AncBE4Max (Addgene plasmid #112094^43^) were a gift from David Liu (Broad Institute). PiggyBac cargo plasmids used in **Figures 3** and **5** and pegRNA-expressing plasmids used in **Figure 4** were created using the Mammalian Toolkit (Addgene article #28197510^44^), which was a gift from Hana El-Samad (UCSF). The sequences of TetR and LacI (Addgene plasmid #81253^16^) were a gift from Timothy Lu. Dox- and IPTG-inducible U6 promoters were initially adopted from the same publication^16^. The IPTG inducible U6 promoter included in that publication consisted of one LacO1 and one LacOS sequence in the promoter and an additional LacOS sequence downstream of the guide RNA. We initially used LacO1 and LacOS in the promoter, but eventually used a promoter with two LacOS sequences in the promoter. In the chemical inducer recording experiment, we also used constitutive U6 promoters with mutations we added in an attempt to reduce their expression. All plasmids are listed in **Supplementary Table 4** and annotated full plasmid maps are available at github.com/liusynevolab/peCHYRON. Pools of PiggyBac cargo plasmids that can be used to make peCHYRON cell lines will be made available from Addgene.

### Cell culture and transfection

293T (CRL-3216) and WT SV40-immortalized mouse embryonic fibroblast (iMEF, CRL-2907) cells were purchased from ATCC, at which point they were certified mycoplasma-free. Cells were grown in DMEM, high glucose, GlutaMAX^TM^ Supplement (Gibco #10566024), supplemented with 10% FBS (Sigma #12306C), at 37°C and 5% CO_2_.

293T cells were transfected with Fugene (Promega #E2311), using a ratio of 1 μg DNA to 3 μL Fugene mixed together in serum-free DMEM or OptiMEM (Gibco #31985062). Unless otherwise noted, the target site for mutation was the genomic HEKsite3. The sequence of this site can be found in **Supplementary Table 4**, in the list of reference sequences for each sequencing experiment. iMEFs were transfected with Lipofectamine 3000 (Invitrogen #L3000015), using a ratio of 1 μg DNA to 1 μL P3000 reagent to 4μL Lipo3000 reagent, mixed together in OptiMEM.

To create the 293T-peCHYRON_6xHis_ cell lines, 293T cells were transfected with plasmids expressing PE2 and a pegRNA that directs the installation of a 20-bp insertion based on the 6xHis sequence, then single colonies were isolated in two rounds of colony picking, dilution of isolated cells, and plating. Integration into the targeted genomic locus and how many of the three copies of this chromosome were modified were verified by deep sequencing. Which clone of these cell lines was used in each experiment is noted in **Supplementary Table 4**.

For transient transfections in 6-well dishes, 1.5 μg prime editor-expressing plasmid was mixed with 500 ng pegRNAs. To create cell lines using PiggyBac integration, cargo mixes (detailed in **Supplementary Table 4**) were mixed with 1/10 – 1/20 total plasmid mass of Super PiggyBac Transposase Expression Vector (System Bioscience, Inc. #PB210PA-1). Stable integrants were selected with 1-2 μg/mL puromycin, 10 μg/mL blasticidin, or 700 μg/mL hygromycin. For the chemical induction experiments shown in **Figure 5** and **Extended Data Figures 8-10**, cells were treated with 2 μg/mL dox or 2 mM IPTG where indicated.

### Flow cytometry

Cell sorting for the experiment shown in **Figure 3** was performed on a Sony MA900 with 4 excitation lasers (488 nm, 638 nm, 405 nm, and 561 nm). Cell sorting for the experiment shown in **Extended Data Figure 4** was performed on the same instrument, and flow cytometry for this experiment was performed on this instrument or on a Sony SH800S with the same lasers. For that experiment, compensation for sfGFP was performed using a stable cell line that expressed only the B→A pegRNA-sfGFP construct. Compensation for mCherry was performed using an equivalent stable cell line expressing only the A→B pegRNA-mCherry construct. Compensation for mTagBFP2 was performed using 293T cells transiently transfected with mTagBFP2. Analysis was performed with software available at floreada.io. FCS files are available at github.com/liusynevolab/peCHYRON.

### Screening pegRNAs for efficient prime editing

Unless stated otherwise, experiments toward finding efficient parts for peCHYRON utilized transient transfections of constructs encoding the indicated components with 3 days incubation before ending the experiment by freezing cell pellets. This pertains to **Figure 2a-d** and **f**, **Extended Data Figure 1a-b** and **d-e**, and **Extended Data Figure 2a**. For the experiment shown in **Extended Data Figure 2b**, plasmids encoding PE2 and a pegRNA installing each B sequence were transfected into 293T_6xHis_ cells in two rounds over 8 days. Then, a plasmid encoding PE2 and libraries encoding the corresponding 6xHis-inserting pegRNAs were transfected into each sample, and cells were collected and analyzed after 2 days.

### Lineage tracing assays

For the reconstruction shown in **Figure 4** and **Extended Data Figure 6**, 293T-peCHYRON_6xHis_ cells in a 6-well dish were transfected with 1.5 μg pCMV-PE2 and 0.5 μg of a mix of equal amounts of 42 pegRNA-expression vectors (**Supplementary Table 4**). These cells were allowed to grow for 8 days, passaging once. Then (“day one”), 20,000 of these cells were plated in each of 4 wells of a 96-well plate, then transfected the next day with the same mix of plasmids. On day four, when they had expanded to approximately 135,000 cells per well, the cells were trypsinized and each well was split into 2 wells of a 24-well dish, before re-transfecting on day five. On day seven, the cells were trypsinized and the entire contents of each well were moved to one well of a 12-well dish. On day nine, when cells had expanded to approximately 750,000 cells per well, each well was split into two wells of a 6-well dish. On day eleven, when each well had expanded to approximately 1.6 million cells, all wells were collected and analyzed by amplicon sequencing.

### Illumina sequencing library preparation

Genomic DNA was isolated with a QIAamp DNA Mini Kit (Qiagen #51304). After DNA extraction, the appropriate region was amplified by PCR. The primers contained partial Illumina adapters and a 5 – 7 nt sample-specific barcode (**Supplementary Table 4**). The PCR was performed with Phusion HotStart Flex DNA Polymerase with GC buffer, with or without DMSO (NEB), 30-35 cycles.

For the experiments shown in **Figure 2a-b**, **Extended Data Figure 1a** and d, and **Extended Data Figure 2c**, reactions were pooled and purified by binding to AMPure beads (0.9 beads:1 sample). The libraries were sent to Genewiz, Inc. for Amplicon-EZ sequencing, where they were further amplified to incorporate the TruSeq HT i5 and i7 adaptors and then sequenced on an Illumina HiSeq 2500 with a paired-end 250 protocol.

For the experiments shown in **Figure 2c**, **Extended Data Figure 2a-b**, **Extended Data Figure 3a**, and **Figure 4**, reactions were purified individually by binding to AMPure beads (0.9 beads:1 sample). Then, they were further amplified to incorporate the TruSeq HT i5 and i7 adaptors, using Q5 High Fidelity DNA Polymerase with GC enhancer, for 15 cycles. The amplified libraries were pooled and purified again by binding to AMPure XP (Beckman-Coulter #A63880) beads (0.9 beads:1 sample). They were subsequently sequenced on an Illumina HiSeq using a paired-end 150-bp protocol at Novogene, Inc. Experiments shown in **Figure 5** were prepared as above, except they were not purified before reamplification and were sequenced on an Illumina NovaSeq X using a paired-end 150-bp protocol at Novogene, Inc. The experiment shown in **Figure 3c-e** and **Extended Data Figure 5d** were prepared as above, except they were not purified before reamplification and were sequenced on an Illumina MiniSeq using a paired-end 150-bp protocol. Experiments shown **in Extended Data Figure 7** were prepared as above, except they were not purified before reamplification and were sequenced on an Illumina MiSeq using a paired-end 300-bp protocol by the UCI Genomics Research and Technology Hub.

### Nanopore sequencing library preparation

For the experiments shown in **Extended Data Figure 1b** and e, **Extended Data Figure 4a-b** and **e-f**, **Figure 3a-b** and **f**, and **Extended Data Figure 5a-d**, genomic DNA was isolated with a QIAamp DNA Mini Kit (Qiagen #51304). After DNA extraction, the appropriate region was amplified by PCR. The primers contained a 10-nt sample-specific barcode (**Supplementary Table 4**). The PCRs were performed with Phusion HotStart Flex DNA Polymerase with GC buffer, with or without DMSO (NEB), 30-35 cycles. The sequences then underwent end repair, A-tailing, and ligation of Oxford Nanopore adaptors using Oxford Nanopore Ligation Kit 14 per the manufacturer’s instructions, except all reagent volumes were divided in half. Libraries were then sequenced on an appropriate MinION flow cell (R10.4.1) and basecalled using Oxford Nanopore’s standard software (Guppy or Dorado).

### High throughput sequencing analysis

The sequences retrieved by HTS were first demultiplexed based on their barcodes; for Illumina sequencing, the script is available at github.com/liusynevolab/CHYRON-NGS. For nanopore sequencing, sequences were demultiplexed using the Maple pipeline, available at https://github.com/gordonrix/maple. Maple outputs demultiplexed .bam files, which we converted to .fastq using the bedtools command bamtofastq. For the experiments shown in **Extended Data Figure 2a-b**, data were analyzed as in^14^. Scripts are available at github.com/liusynevolab/CHYRON-NGS. For the experiments shown in **Figure 2a-b**; **Extended Data Figure 1a** and d; BE samples in **Extended Data Figure 1b**, and **Extended Data Figure 2c**, data were analyzed with CRISPResso2^45^. For the high-throughput pegRNA screen shown in **Figure 2c**, 20-bp insertion sequences were extracted from fastq files by simply grabbing the 20-letter sequence at the expected edit site, then comparing it to the wt (unedited) sequence. If the extracted sequence exceeded a Hamming Distance cutoff from the unedited sequence, it was considered a real insertion for further analysis. All insertion sequences were tabulated, and instances of identical insertions were tallied. To calculate enrichment scores, Illumina sequencing reads from the pegRNA hp-miniprep pool (the plasmids that were transfected into 293T_6xHis_ cells at the start of the experiment) were analyzed in the same manner. The abundance of each RT template sequence in the original pegRNA pool was tabulated. Finally, the enrichment of each 20-bp insertion sequence was calculated as follows, where *g* is the proportion of genomic reads bearing that insertion sequence, and *m* is the proportion of library plasmid (the hp-miniprep pool) reads bearing that insertion sequence:

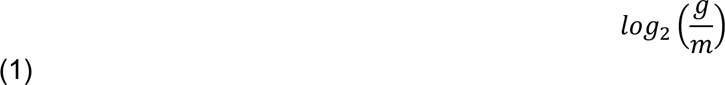

For **Extended Data Figure 3a** and **Figure 4**, for each sample, associated forward and reverse reads were merged (PEAR 0.9.10^46^). For **Figure 3c-e**, **Figure 5**, and **Extended Data Figure 7**, forward reads were used. For **Extended Data Figure 1d-e**, **Extended Data Figure 4a-b** and **e-f**, **Figure 3a-b** and **f**, and **Extended Data Figure 5a-d**, nanopore sequences were used. Insertion sequences were extracted by finding the expected sequence motif upstream of the edit site, then searching for the expected sequence motif downstream of the edit site. For each read, the insertion length was increased in increments of 20 bp until the expected sequence motif downstream of the edit site was found. The full insertion sequence was extracted and the 17-bp propagator sequences in each insertion were compared to the known propagator sequences to ensure only legitimate insertions were grabbed. The 3-bp signature mutations in each insertion were converted to 1-character symbols to improve the ease of interpreting results. Insertion sequences represented by 1-character signature mutations were tabulated, and the counts of each insertion sequence were tallied. In parallel, the lengths of insertion sequences between the upstream motif and downstream motif were tallied, even if they were not multiples of 20 and/or did not contain the expected propagator sequences.

All the analyses were done in Python. The scripts and detailed instructions are available at github.com/liusynevolab/peCHYRON unless otherwise specified.

### Calculation of cannibalization

For **Extended Data Figure 4a-b** and **Figure 3a**, nanopore sequencing of pegRNA genes was used to characterize gene length over time – the longer read lengths of which nanopore is capable allowed us to use amplicons >1 kb whose efficiency would not be biased by the insertions or deletions we are trying to detect. Because nanopore sequencing produces a significant rate of insertion and deletion errors, pegRNA gene sequences were compared to sequences of the original pegRNA gene plasmid, amplified and library-prepped under the same conditions, sequenced on the same platform, basecalled with the same software version, and analyzed with the same script version. After demultiplexing as described above, the sizes of all sequences that lie between a motif at the beginning of the pegRNA coding sequence and a motif downstream of the coding sequence and protective nick site was computed. The range of sizes that included 98% of reads for the plasmid was defined as the “unchanged” size. Then the proportion of sequences that were shorter or longer than the unchanged size was computed for each sample, and the proportion of such sequences in the plasmid sequence was subtracted from the value.

### Lineage reconstruction

To reconstruct cell lineage in **Figure 4**, we created a list of all insertion sequences in each of the 13 wells used for the analysis. Each insertion has an abundance, based on the number of HTS reads that include that exact insertion sequence, and a length, equal to the number of peCHYRON insertions at the recording site. For our initial analysis, the researcher performing the analysis (TBL) was not told which well was which. We refined the list for each well to include only those insertions that passed two filtering steps: 1) a length filter that excluded any sequences with 3 or fewer inserted sequences and 2) an abundance filter that removed everything after a decreasing convex knee found using the kneedle algorithm^29^. For each sequence in this set, we generated all possible subsequences containing the first insertion (*i.e.* prefixes) and assigned a weight equal to the inverse of the number of prefixes. This results in a multiset of prefixes for each sequence. Afterwards, the distances between the multisets were found using the generalized Jaccard distance for multisets. We used this set of distances to reconstruct the relationships using the UPGMA^47^ hierarchical clustering algorithm. (github.com/scipy/scipy/blob/v1.2.1/scipy/cluster/hierarchy.py#L411-L490). The two algorithms (Jaccard and Prefix Jaccard) were compared by computing the Robinson Foulds score^48^, a metric that compares a reconstruction to a ground-truth tree, for each reconstruction (**Extended Data Figure 6b**). For each percentage of downsampling between 99 and 5%, 1000 different data downsampling operations were performed. Robinson Foulds scores were computed for each using the ete3 package^49^ (http://etetoolkit.org/).

All the analyses were done in Python. The scripts and detailed instructions are available at github.com/liusynevolab/peCHYRON.

### Event reconstruction via bigram frequencies

To evaluate bigram frequencies of signatures at recording loci, ‘expanded bigrams’ were defined as any ordered pair of signatures in a record, taking into account non-contiguous pairs. In other words, a single signature produces a bigram for each signature after it in a record (*e.g.*, for a four-signature record ABCD, the expanded bigrams produced from signature A are AB, AC, and AD; the expanded bigrams of signature B are BC and BD). Consecutive unigrams of the same signature were treated as a single unigram (*e.g.* AAAB is treated as AB) and reduced from multiple to single signatures where applicable (referred to as ‘bigram reduction’). The read counts of all bigrams were enumerated by sample to assess the total frequency of each bigram across all insertions. For transfection epoch experiments in **Extended Data Figure 7A**, after removal of signature sequences not included in the overall experiment (the result of sequencing error), the total bigram frequencies were plotted as a heatmap with the signature in position 1 (‘P1’) and position 2 (‘P2’).

### Event reconstruction via stretched records

Samples were sequenced with Illumina. The resulting fastq files were aligned to a reference sequence for the peCHYRON recording locus, and insertions at the recording locus were extracted. The insertions were analyzed in 20 bp increments, where the first 3 bp are the signature mutation and the last 17 bp are the propagator sequence. If propagator sequences didn’t match their expected sequences within a hamming distance of 2, they were assumed to be incorrect amplicons from sequencing library prep and were thrown out. If the insertions passed this first filter, the 3 bp signature mutations were extracted and converted to 1-letter symbols according to a user-defined signature key (this is to simplify interpretability of records). Then, the resulting records were filtered again to remove reads with unexpected signatures due to sequencing error (for example, samples from the dox and IPTG recording experiment should only have signatures corresponding to the dox- or IPTG-inducible pegRNAs or the constitutive pegRNA). This final list of records was computationally “stretched” to the least common multiple of the lengths of all records. Insertion lengths with fewer than 100 reads were excluded from the analysis. To assign weights to records of different lengths, the number of reads corresponding to each insertion length was first tallied. Then each insertion length was given a weight as described in equation 2.

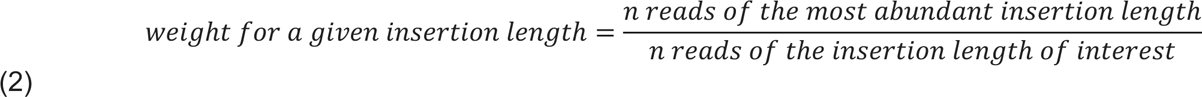

For example, if records with 2 insertions were most abundant, and there were 100,000 records with 2 insertions but only 20,000 records with 3 insertions, the weight for records with 3 insertions would be 5 (100,000 / 20,000). When calculating the fraction of each signature at each insertion length, records were pooled together according to their weight (*e.g.*, a record given a weight of 5 was added to the pool 5 times). The fraction of each signature is simply the % of records in the pool with a particular signature at each position.

For the samples exposed to chemical epochs (**Figure 5c** and **Extended Data Figure 10**), all records were first stretched to a length of 60, because this was the least common multiple of the experimental insertion lengths. Then, instead of plotting the fraction of each signature at each position, the fold-change relative to the uninduced case was plotted. The fraction of each signature in the uninduced case was calculated from a control sample that was never exposed to any inducer for 18 days. To calculate fold-change, the experimental samples’ fractions of each signature at each position were first computed, then divided by the fractions observed in the uninduced control sample. The Spearman rank coefficients between the stretched records of pairs of samples were computed as follows: for each of dox and IPTG, the signature proportions across all 60 points of the stretched array were ranked and plotted in each of X and Y dimensions for the two sample pairs being compared and the correlation coefficient determined. The average of D and I coefficients was calculated to obtain a single value for comparison. The resulting 27x27 coefficient matrix was hierarchically clustered using the single linkage method in the Scipy.cluster.hierachial module in the SciPy library.

All these analyses were conducted in Python. Refer to github.com/liusynevolab/peCHYRON for scripts and detailed instructions for use.

### K-means clustering for event reconstruction

The total frequency of each signature across all insertions were summed for each sample (without ‘bigram’ reduction) and univariate K-means clustering was performed to group samples together based upon total IPTG and total dox signature frequency. Clustered data was plotted in **Extended Data Figure 8** as barplots of fraction of total signatures which are D or I, with clustering visualized by bar color, with each replicate clustered and plotted separately. For chemical epoch experiments in **Extended Data Figure 9**, total bigram frequencies were divided by frequencies of the bigram’s anadrome (e.g. DI frequency divided by ID frequency), and then log_2_ transformed to obtain the log_2_ of the fraction of a bigram vs its anadrome. The log_2_ ratios of bigram/anadrome pairs were filtered to only samples which clustered into dox-positive and/or IPTG-positive groups from analysis of total signatures per sample. The filtered data was then subjected to univariate K-means clustering, grouping samples into clusters based upon their expected order of induction, with each replicate clustered and plotted separately.

Data was analyzed and visualized using Python3. K-means clustering was performed using the KMeans() function of the sklearn.cluster library, with appropriate cluster number identified with the find_peaks() function of the scipy.signal library. Barplots were prepared using the barplot() function of the seaborn library.

### Entropy calculations

To calculate Shannon entropy^28^, we first made a table of all the signature sequences in the relevant dataset with, for each sequence, the number of times it was observed (the “count”). We did one calculation for signatures inserted at “odd” positions (1, 3, 5, etc.) and one for “even” signatures. Once a table of signatures and counts (c) was created, we calculated the proportion (*p*) for each sequence (equation shown here for sequence i).

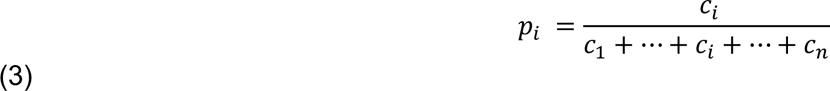

Then we calculated the overall Shannon entropy (H) for the dataset:

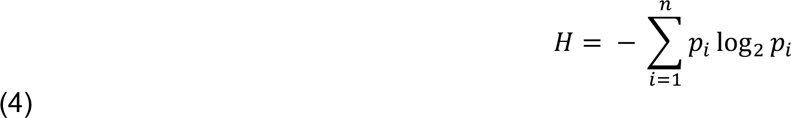

All analyses were done in Excel (**Supplementary Table 2**).

### Calculation of signature repetition

For the analysis shown in Figure 3e, the number of repeated signatures in each 6-signature read was calculated by hand. First, we calculated the number of repeats of signatures installed by A→B pegRNAs, found in positions 1, 3, and 5. Then we calculated the repeated signatures installed by B→A pegRNAs, found in positions 2, 4, and 6. The maximum number of repeats was 4, found in sequences in which the signatures in positions 3 and 5 matched position 1, and the signatures in positions 4 and 6 matched position 2.

### Statistical analyses

In all cases, biological replicates were derived from different populations of cells that were manipulated separately throughout the experiment. For technical replicates, cells were grown and manipulated, and DNA extracted, together. All procedures downstream of DNA extraction were performed separately. Significance of the results in **Figure 2d** was calculated in GraphPad Prism by Welch’s t test. Significance of the results in **Extended Data Figure 4a** was calculated in GraphPad Prism by unpaired t test without Welch’s correction.

### Figure preparation

The **Graphical Abstract,** Figure 1**, Extended Data Figure 1c, Extended Data Figure 3b**, a portion of Figure 4a, and **Extended Data Figure 7a-b** were prepared with InkScape. Figure 2c, **Extended Data Figure 7d**, and **Extended Data Figure 10** were plotted in Excel, then converted to scalable vector graphics with InkScape. A portion of Figure 4a was prepared at biorender.com. The plots in Figure 4b-c and **Extended Data Figure 6a** were generated using the hierarchy.dendrogram function in matplotlib (scipy.org). Heatmaps of transfection epoch bigram frequencies in **Extended Data Figure 7c** were plotted using the heatmap() function of the seaborn library in Python3. For **Extended Data Figures 8-9**, scatterplots and barplots were prepared using the scatterplot() and barplot() functions of the seaborn library, respectively. All other plots were created in GraphPad Prism.

## Data Availability Statement

All scripts created for this manuscript are available at github.com/liusynevolab/peCHYRON. All NGS data sets will be deposited at the NCBI’s Sequence Read Archive upon publication, and can be accessed with the keyword “peCHYRON.” Full plasmid maps are available at github.com/liusynevolab/peCHYRON. Plasmids necessary to carry out peCHYRON lineage tracing will be available at Addgene. See **Supplementary Table 4** for a guide to these data and reagents. Please contact CCL and TBL for other reagents.

---

**Extended Data Figure 1.**
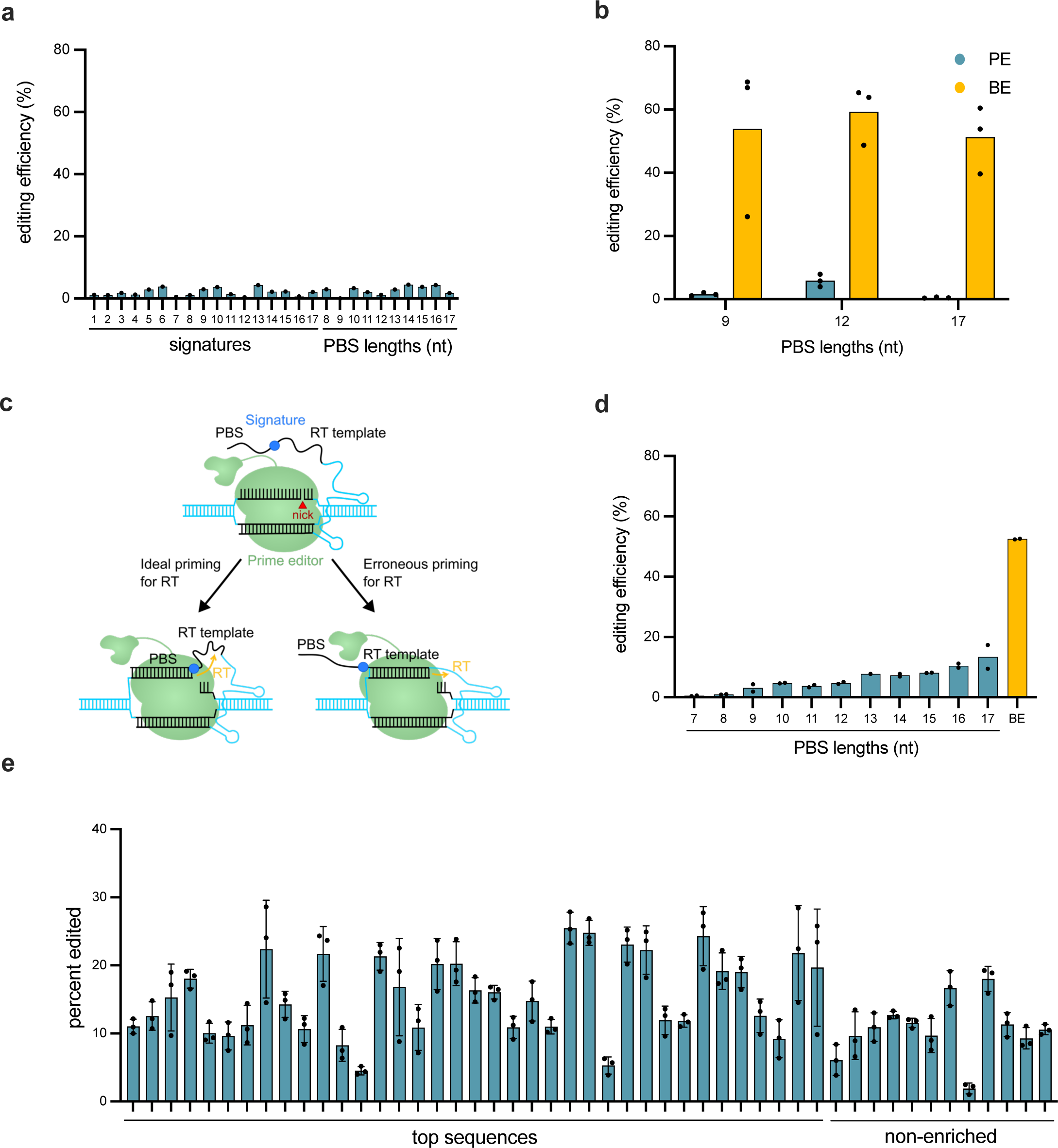
Testing alternative pegRNA candidates for propagation. **a**, Efficiency of iterative editing with a pegRNA that can both target and insert the same 20-bp sequence (A→A) was low. The 20-bp sequence was derived from the 6xHis sequence. Constructs expressing PE2 and appropriate pegRNAs were transfected into 293T_6xHis_ cells. Libraries of all possible signature mutations, each with a different PBS length, were scanned. **b**, Although the efficiency of inserting the 6xHis sequence at a 6xHis target site was low, the 6xHis target site readily recruited Cas9. To ensure that the 6xHis sequence could be efficiently targeted by a pegRNA-directed Cas9, plasmids encoding selected pegRNAs from (a) and either PE2 or the C→T base editor AncBE4Max were transfected into 293T_6xHis_ cells, and the efficiency of a prime edit or base edit at the 6xHis sequence was measured. Points represent biological replicates. **c**, Proposed mechanism of failed prime editing with a pegRNA that both targets and inserts the same 20-bp sequence (A→A). The PBS and RT template sequence share identity, so priming can occur erroneously in the RT template instead of the PBS, yielding no net insertion. **d**, The efficiency of inserting the wt site3 sequence at a 6xHis target site was low. Here, site3 was the original sequence at the recording locus and 6xHis had been inserted (A→B) to make the 293T_6xHis_ cells used in this experiment. Thus, to achieve propagation, the efficiency of adding back site3 was tested (B→A). For optimization, multiple lengths of the PBS of the site3-inserting pegRNA were tested. Each sample is a library of all possible signatures with the indicated PBS length. To ensure that the 6xHis sequence could be efficiently targeted by Cas9 directed by this type of pegRNA, plasmids encoding the C→T base editor AncBE4Max and a pegRNA that targets 6xHis were co-transfected into 293T_6xHis_ cells, and the efficiency of a base edit in the 6xHis sequence was measured. Points represent biological replicates. **e**, Sequences that can efficiently insert downstream of 6xHis were identified. Top hits and some non-enriched control sequences from the high-throughput screen shown in Figure 2c were individually cloned, and their insertion efficiencies were assayed. All experiments were conducted in 293T_6xHis_ cells with PEmax; we used lower concentrations of plasmids than usual for this transfection, so insertion efficiencies are expected to be lower than in other experiments. Insertion sequences are ordered from left to right from most-enriched to least-enriched in the original screen.

**Extended Data Figure 2.**
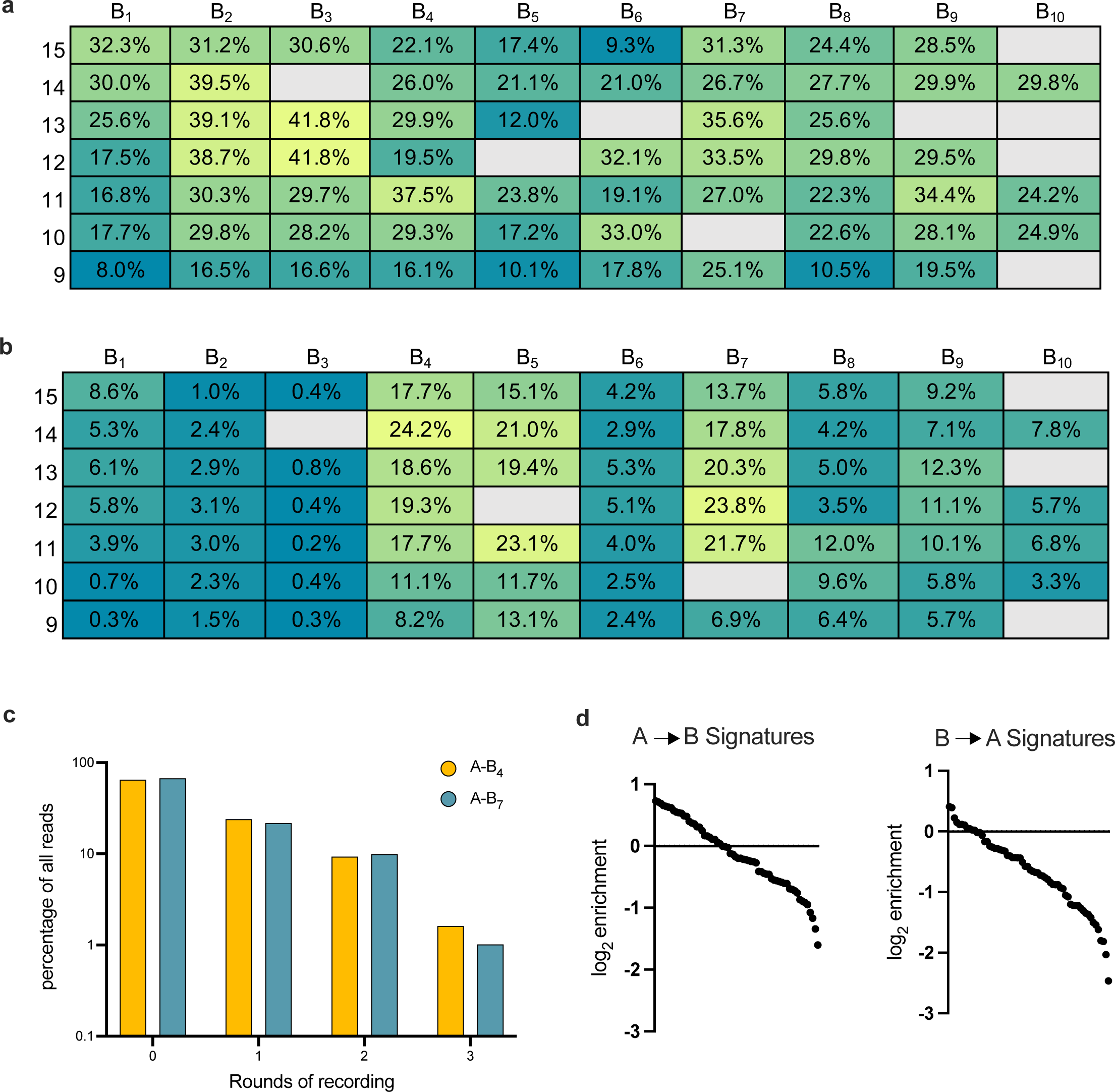
Identification of efficient propagating pegRNAs. **a**, Heatmap of editing efficiencies for the top ten A→B propagator sequences identified in an initial screen similar to Extended Data Figure 1e. For each A→B propagator sequence (B_1_-B_10_), a library of signatures was tested, using PBS lengths ranging from 9-15 nts (row labels). **b**, Heatmap of editing efficiencies for pegRNAs that target the top ten propagator sequences shown in (a) and insert the 20-bp 6xHis sequence (B→A). Libraries encoding pegRNAs with all possible signatures, at each PBS length from 9-15 nts (row labels), were transfected into 293T_6xHis_ cells in which the corresponding B sequence had been installed at the recording locus. **c**, A→B and B→A pegRNAs can achieve propagation. Plasmids expressing PE2 and the top-performing pegRNA libraries identified in (a) and (b) were transfected into 293T_6xHis_ cells. The lengths of insertions at the recording locus were assayed after 2 rounds of transfection over 6 days. Insertions of 20 bp were considered to correspond to 1 round of recording, 40 bp to 2 rounds, and so forth. **d**, A variety of signatures can be installed by the top-performing A-B_4_. libraries of pegRNAs. From the screens shown in (a) and (b), the log2 enrichment of each of the 64 possible signatures was calculated for the A→B (left) and B→A (right) pegRNA libraries.

**Extended Data Figure 3.**
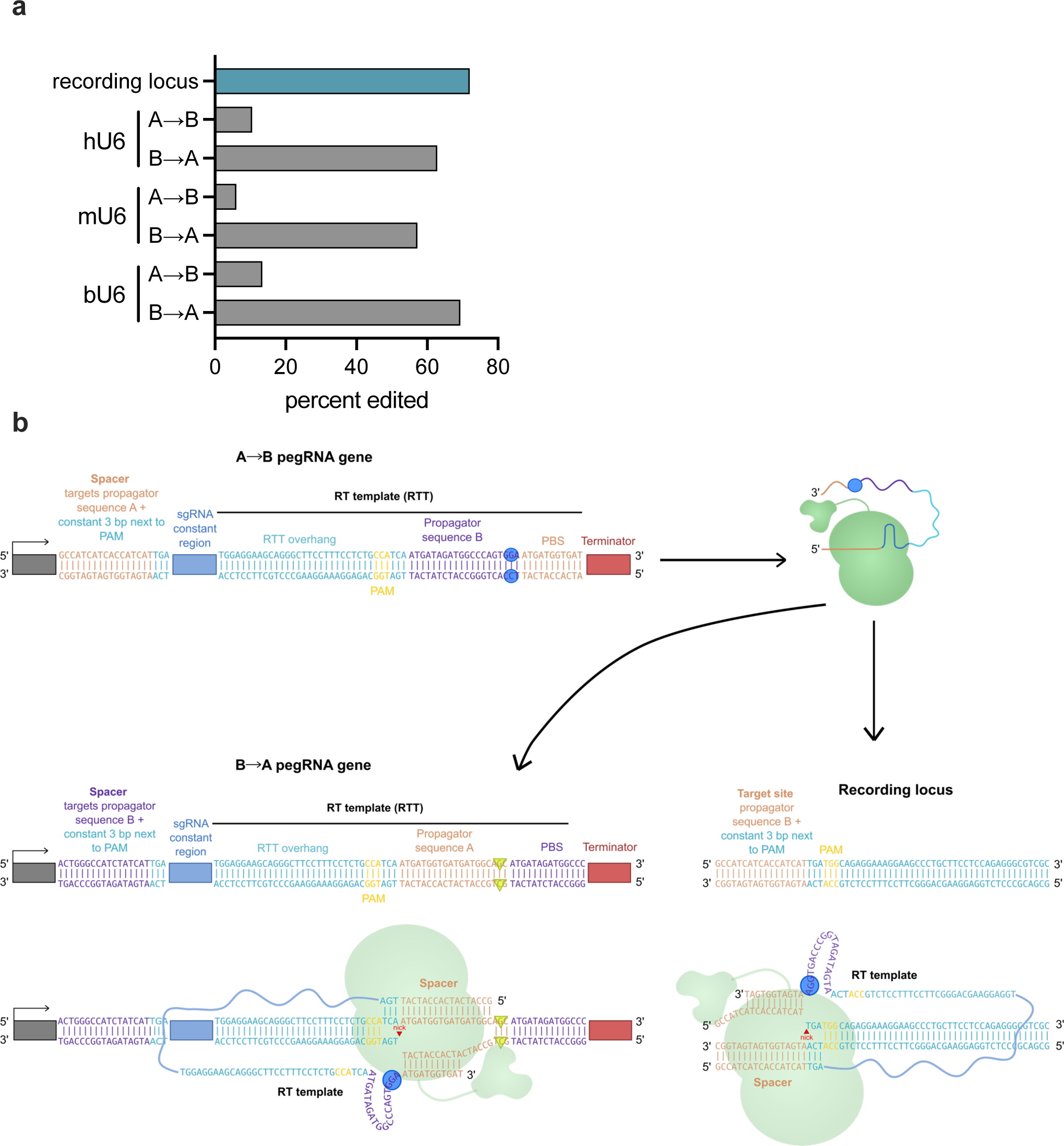
Cannibalization by peCHYRON. **a**, Editing at the recording locus and pegRNA genes in peCHYRON cells. Prime editor and pegRNAs were integrated into the genome of 293T_6xHis_ cells as PiggyBac Transposase cargo, and integrants were selected with puromycin. 45 days after transfection, the recording locus (where editing is desired) and all pegRNA-expressing loci (where cannibalization occurs) were sequenced. The graph indicates the proportion of loci that were edited. Data from loci expressing A→B or B→A pegRNAs from each of three types of U6 promoters, human (h), mouse (m), and bovine (b), are shown. **b**, More detailed cartoon showing cannibalization of B→A pegRNA gene by the A→B pegRNA. After transcription and coupling with prime editor, the spacer sequence of the A→B pegRNA can recognize either the target site at the recording locus or the PAM-adjacent target site on the antisense strand of the B→A pegRNA gene. Note the B→A pegRNA only inserts 17 nt of the target sequence recognized by the A→B pegRNA, but the complete 20-bp target site and PAM are present in the B→A pegRNA gene because prime editing requires a scaffold sequence that complements with the region downstream of the nick, to enable flap equilibration after reverse transcription. This cartoon only shows cannibalization of the B→A pegRNA gene, but cannibalization can also occur in the inverse direction (the A→B pegRNA gene is cannibalized by the B→A pegRNA).

**Extended Data Figure 4.**
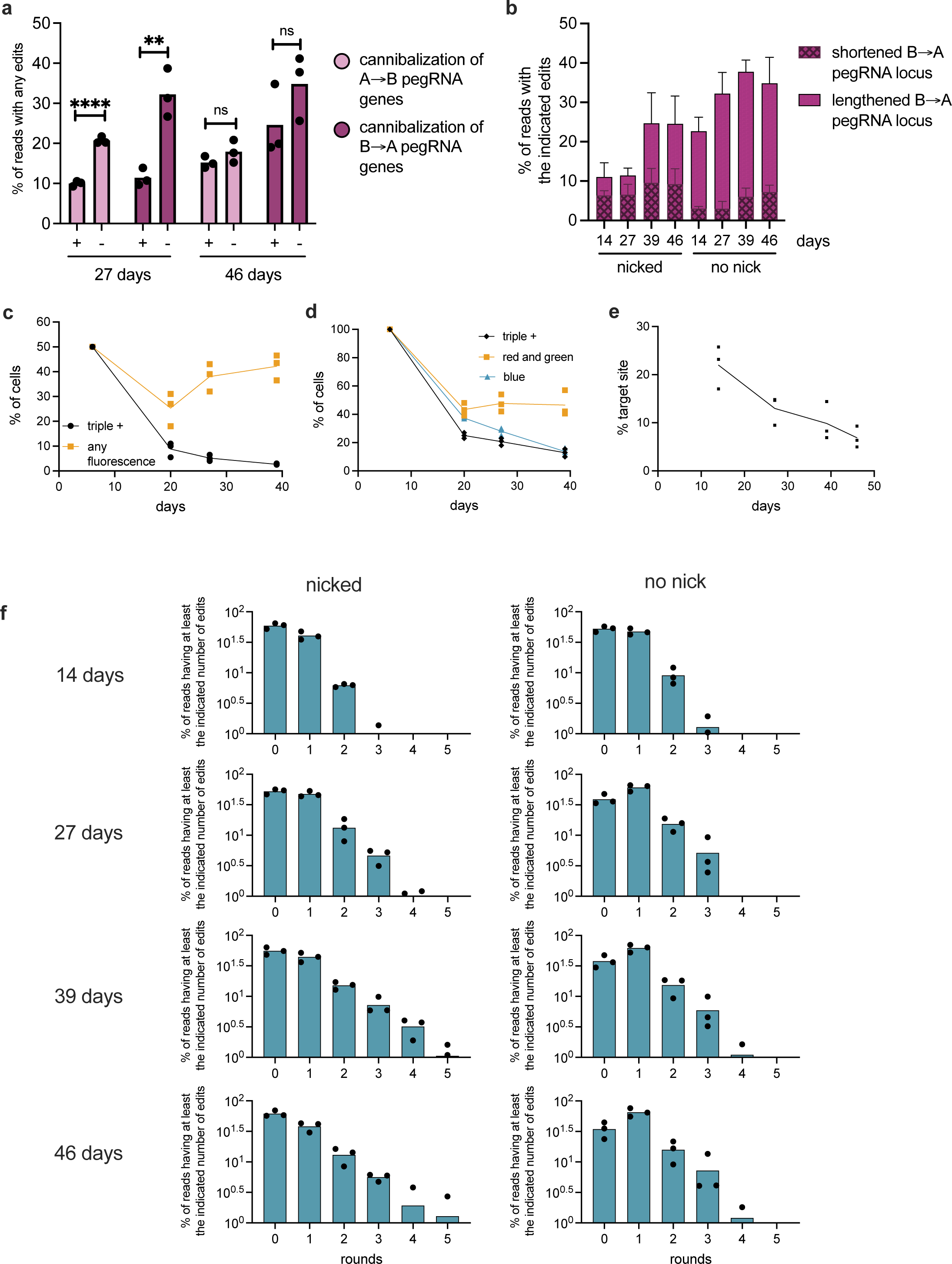
Long-term recording with peCHYRON. **a**, Cannibalization of the peCHYRON system was blocked by a protective nick. Expression cassettes for PEmax (marked with mTagBFP2), A→B (marked with mCherry) and B→A (marked with sfGFP) pegRNAs, and, optionally, protective nicking guides, were co-transfected with PiggyBac transposase for stable integration into the genome of 293T_6xHis_. Six days after transfection, cells positive for blue, red, and green fluorescence were selected and propagated. Cells were collected at the indicated time after the initial transfection, then the A→B and B→A pegRNA-expressing loci were sequenced. Any deviation from the original sequence was considered cannibalization and plotted (see Methods). Throughout this figure, each dot represents one of three biological replicates. At 27 days but not 46 days, the extent of cannibalization with and without the protective nick were significant by unpaired t test (P<0.0001 for A→B and P=0.005 for B→A). **b**, At early timepoints, in the presence of the protective nicking guide, edits. This observation suggests that simultaneous nicking by a cannibalizing pegRNA and by the protective nicking sgRNA can, at a low rate, lead to recognition by the host DNA repair machinery followed by mutagenic repair. Further optimization of the position of the protective nick may alleviate this side effect while continuing to block undesired prime editing. Error bars reflect the standard deviation of 3 biological replicates. **c**, The fitness cost of peCHYRON components was acceptable, but silencing occurred. At the time of initial cell selection, 6 days after transfection, 5,000 cells from each protective-nicking guide replicate were each mixed with 5,000 cells from the parent, non-fluorescent cell line. The proportion of blue, red, and green fluorescent cells was determined by flow cytometry at the indicated time points. The number of cells “triple-positive” for all colors, or positive for any color, is plotted. **d**, In the populations of cells transfected with the protective nicking guide, silencing was tracked by flow cytometry at the indicated timepoints. **e**, Cells tended to lose recording loci over time. In the parent cell line, the first A site is knocked into one copy of the site3 locus, of which there are three copies in 293T cells. When site3 is sequenced by amplicon sequencing, therefore, 20-35% of reads align to the locus including the A site. This proportion was calculated at the timepoints indicated, in the cells containing the protective nicking guide. **f**, For the experiment shown in a-e, the extent of editing at the recording locus in cell lines including (left) or not including (right) the protective nicking guide was calculated. Insertions of 20 bp were considered to correspond to 1 round of insertion, 40 bp to 2 rounds, and so forth. The proportion of loci that had reached at least each number of rounds was plotted, *e.g.*, a locus with an insertion 140-bp long, corresponding to 7 rounds, would be counted towards the total for 1, 2, 3, 4, 5, 6, and 7 rounds.

**Extended Data Figure 5.**
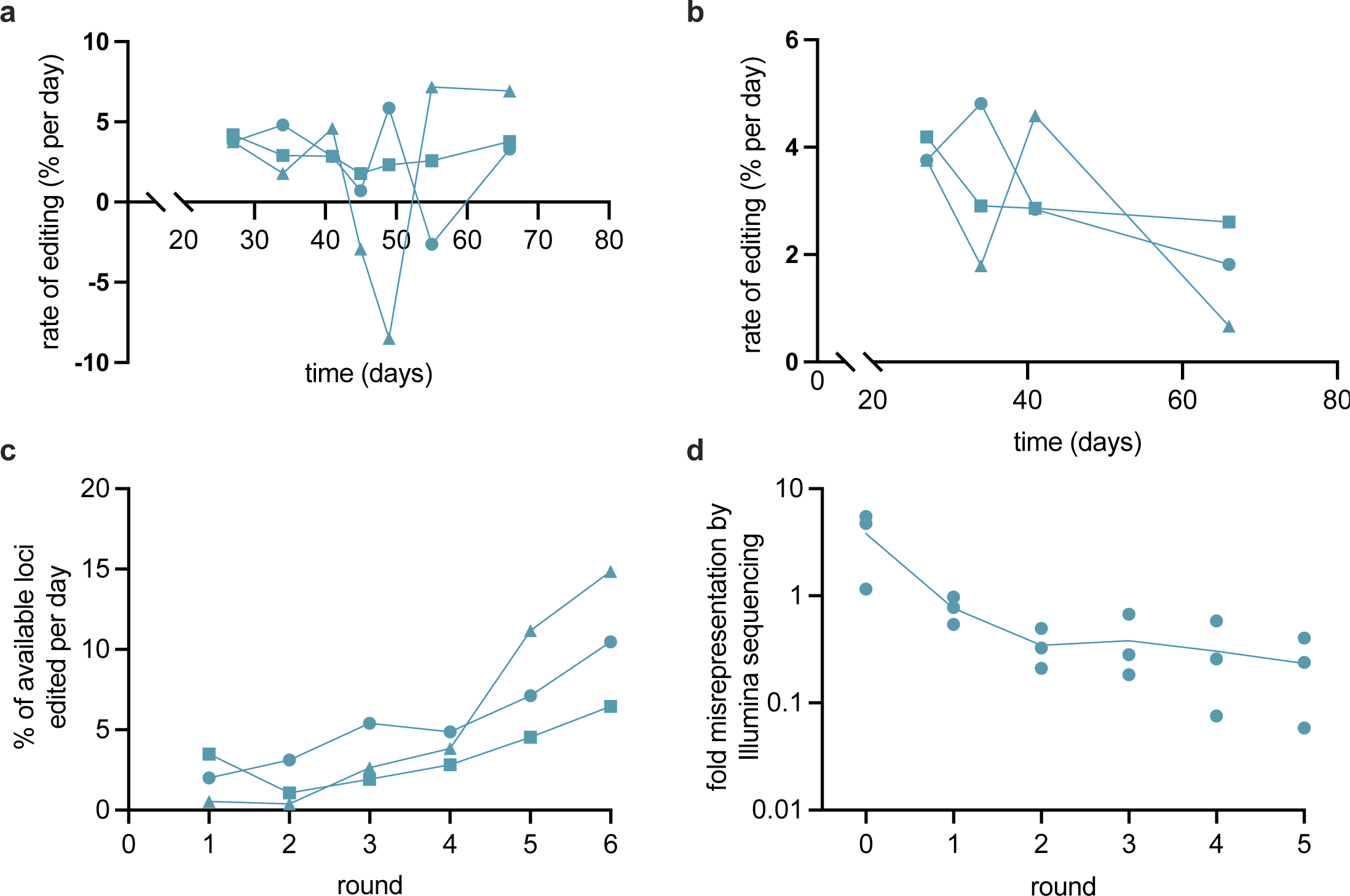
Rates of peCHYRON recording. **a**, Editing rates declined only gradually over the course of the experiment shown in Figure 3a-e. Based on the distribution of edit lengths detected at each timepoint, we calculated the rate of editing between each pair of timepoints as the average change in edits from time N to time N+1. The plotted value at each timepoint is the calculated rate for the period ending at that timepoint. For example, the value at 27 days is the rate from 0-27 days, the value at 34 days is the rate from 27-34 days, *etc.* Each line reflects the editing rates of one biological replicate. Transient negative values likely reflect a slightly lower growth rate in cells that edit faster. **b**, As in a, but with a single rate calculated from 41-66 days. **c**, Rates of editing were maintained as arrays grew longer. For the period from 27 to 34 days, the rate of each conversion step *(e.g.* step 1 converts a locus with no edits to one with a single edit) was calculated for each replicate. **d**, Illumina library prep and sequencing underrepresents longer amplicons. The same samples were sequenced by nanopore after amplification of a 2.3 kb region or by Illumina after amplification of a region that is 154 bp before editing. The percent of reads that showed at least that many rounds of editing by Illumina was divided by the same value obtained from nanopore sequencing.

**Extended Data Figure 6.**
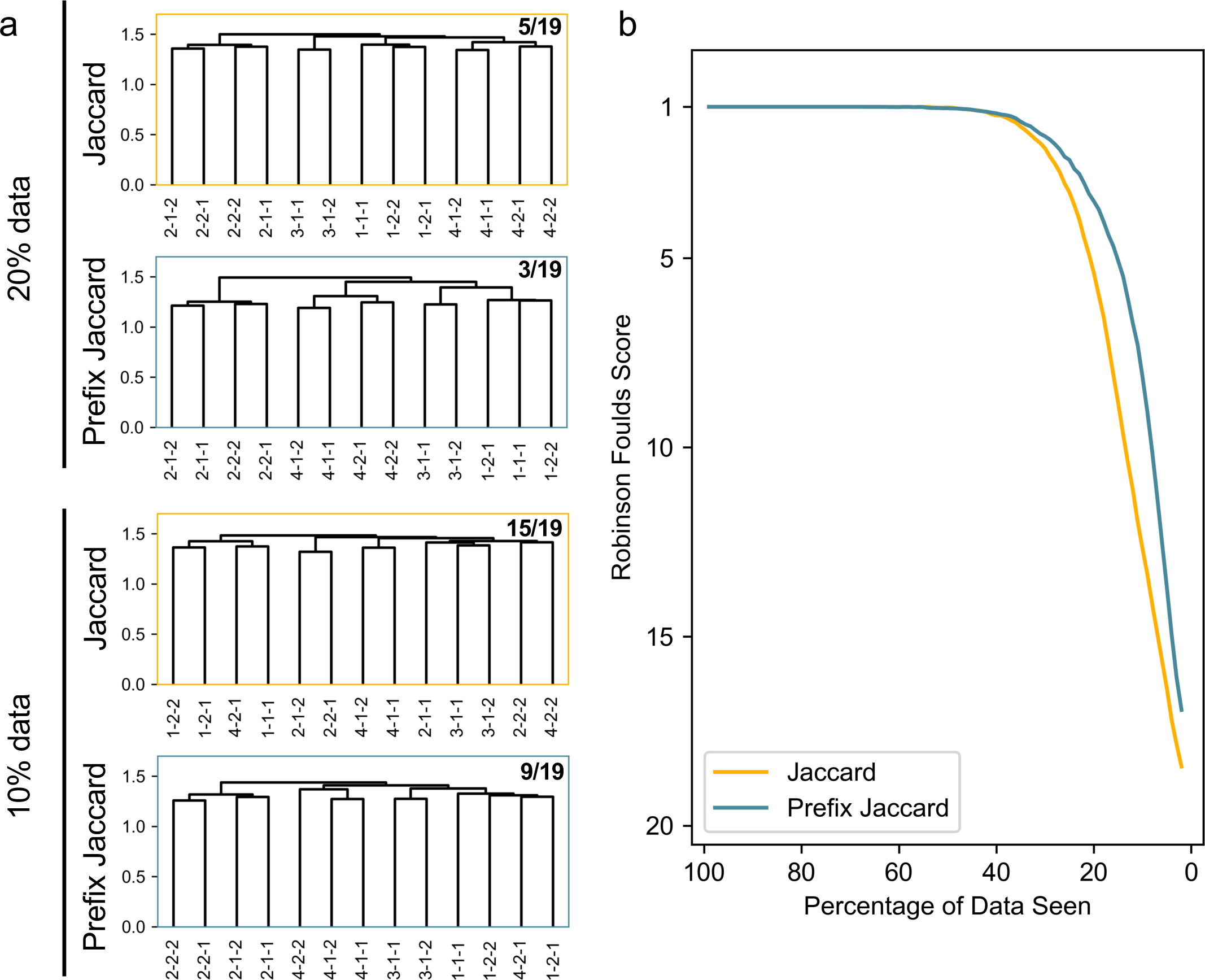
Comparison of lineage reconstruction methods. **a,** Reconstruction of culture splitting patterns from a cell lineage tracing experiment with random downsampling was more accurate when our prefix Jaccard similarity index was used, rather than the traditional Jaccard similarity index. For each reconstruction, we calculated the Robinson Foulds score, which is the number of changes required to transform the reconstruction into the ground truth tree. The score for each reconstruction is shown in the upper right, divided by the highest possible score (19). **b,** Average Robinson Foulds scores for reconstructions using our prefix algorithm (blue) and the traditional Jaccard distance (orange). This was calculated by randomly downsampling the data to a certain percentage (x-axis) and performing a reconstruction which was then compared to the true tree. Each point was obtained by averaging over 1000 repeats. Note, perfect reconstruction in our case returns a score of 1 instead of 0 because we compare a rooted reconstructed tree to an unrooted ground truth tree.

**Extended Data Figure 7.**
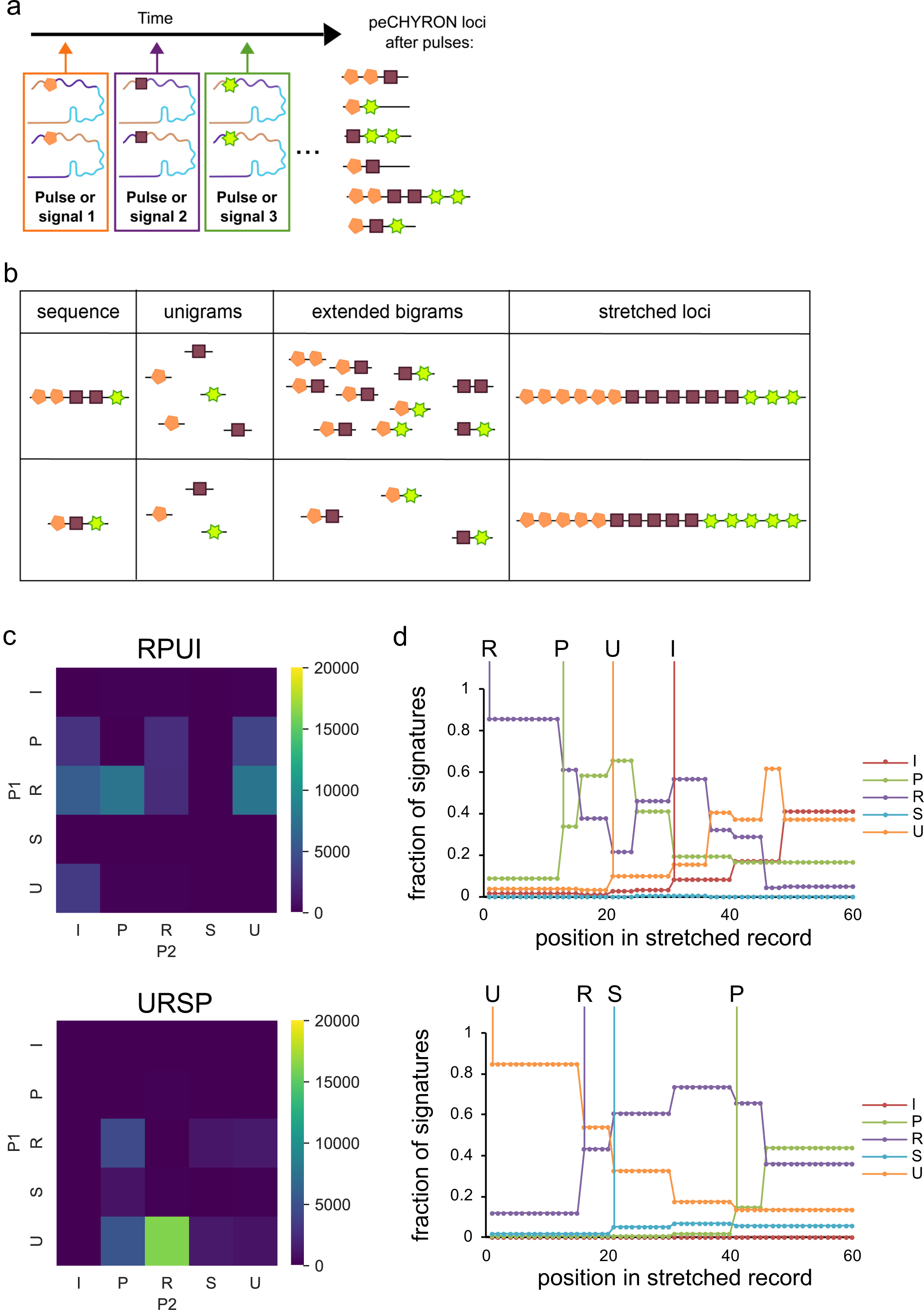
Recording of transfection order with peCHYRON. **a**, General approach to recording pulses (or any time-dependent signals) that are linked to expression of pegRNAs with known signature mutations. A pair of A→B and B→A pegRNAs are expressed together for any signal to be recorded, so that one pegRNA of the pair will record the signal regardless of the state (A or B) of the active link in the recording locus, and both pegRNAs can continuously record the signal if it is present for a long duration. Colored shapes represent signature mutations. **b**, Methods of peCHYRON sequence analysis. For the simplest analysis, which treatments happened in a cell population can be inferred by calculating the proportion of each unigram, or type of signature. The order of treatments can be deciphered by considering the extended bigrams of signature mutations it contains. Essentially, each signatures forms one bigram with every signature that appears after it in a read. Alternatively, records can be analyzed by stretching each record to a constant length, so it may be pooled with all other records from a population of cells and used to calculate the proportion of each signature at each position along the stretched records. **c**, Extended bigrams of recorded signatures reflected the order of transfections. 293T_6xHis_ cells, in which PEmax and the protective nicking sgRNA were stably expressed, were transfected with pairs of pegRNAs during 4 epochs of 3-6 days each. After the 4^th^ epoch, recording loci were sequenced. Heatmaps show the relative abundance of all possible ordered pairs; 3-letter signatures are represented by a single letter. The true epoch pattern is above each heat map. P1, first signature in a bigram. P2, second signature in a bigram. **d**, Computational stretching can also reveal the order of transfections. For the experiment also shown in c, the signature mutations were extracted from peCHYRON records, then the sequence of signature mutations from each recording locus was computationally stretched to a constant length (this is 60 because it is the least common multiple of all insertion lengths in the data set). Stretched records were pooled together in a manner that allows each insertion length to contribute equally to the final pool long insertions are weighted more heavily to prevent being overshadowed by short, abundant insertions that are less informationally rich (*i.e.*, records of each length were weighted by the inverse of their abundance in the total pool). At every position along the stretched records, the proportions of each signature were plotted. Each vertical line and signature label corresponds to the first increase in the abundance of the corresponding signature.

**Extended Data Figure 8.**
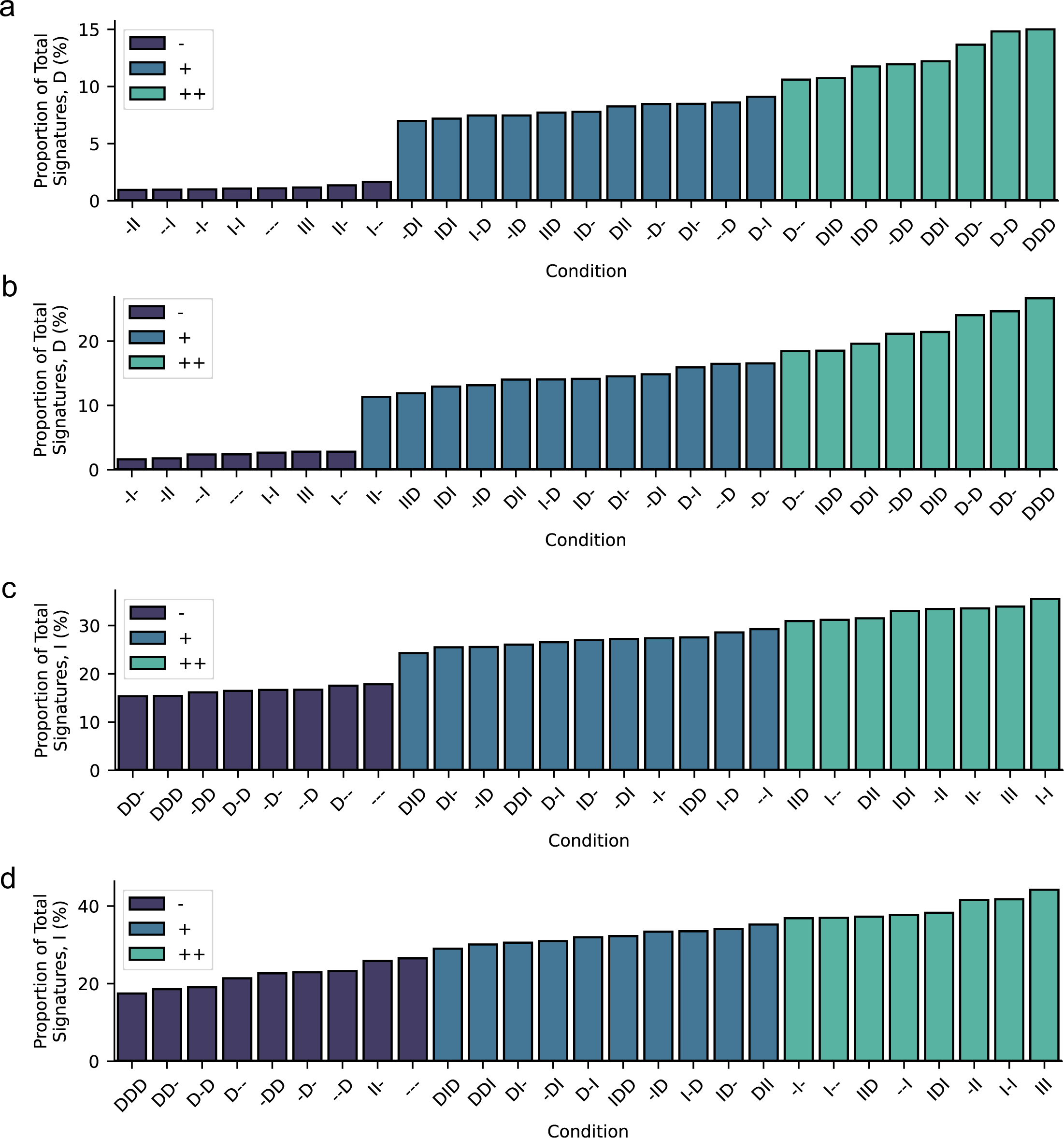
Recording of inducer duration with peCHYRON. **a**, As described for Figure 5a, the proportions of signatures in Replicate 1 samples that were D were calculated. We then applied k-means clustering on these values, choosing the cluster number by automatic evaluation of an elbow plot by identifying the number of clusters for which the second derivative of the inertia is a peak; bars are colored based on cluster membership. **b**, As in a, for Replicate 2. **c**, As in a, but for I. **d**, As in c, for Replicate 2.

**Extended Data Figure 9.**
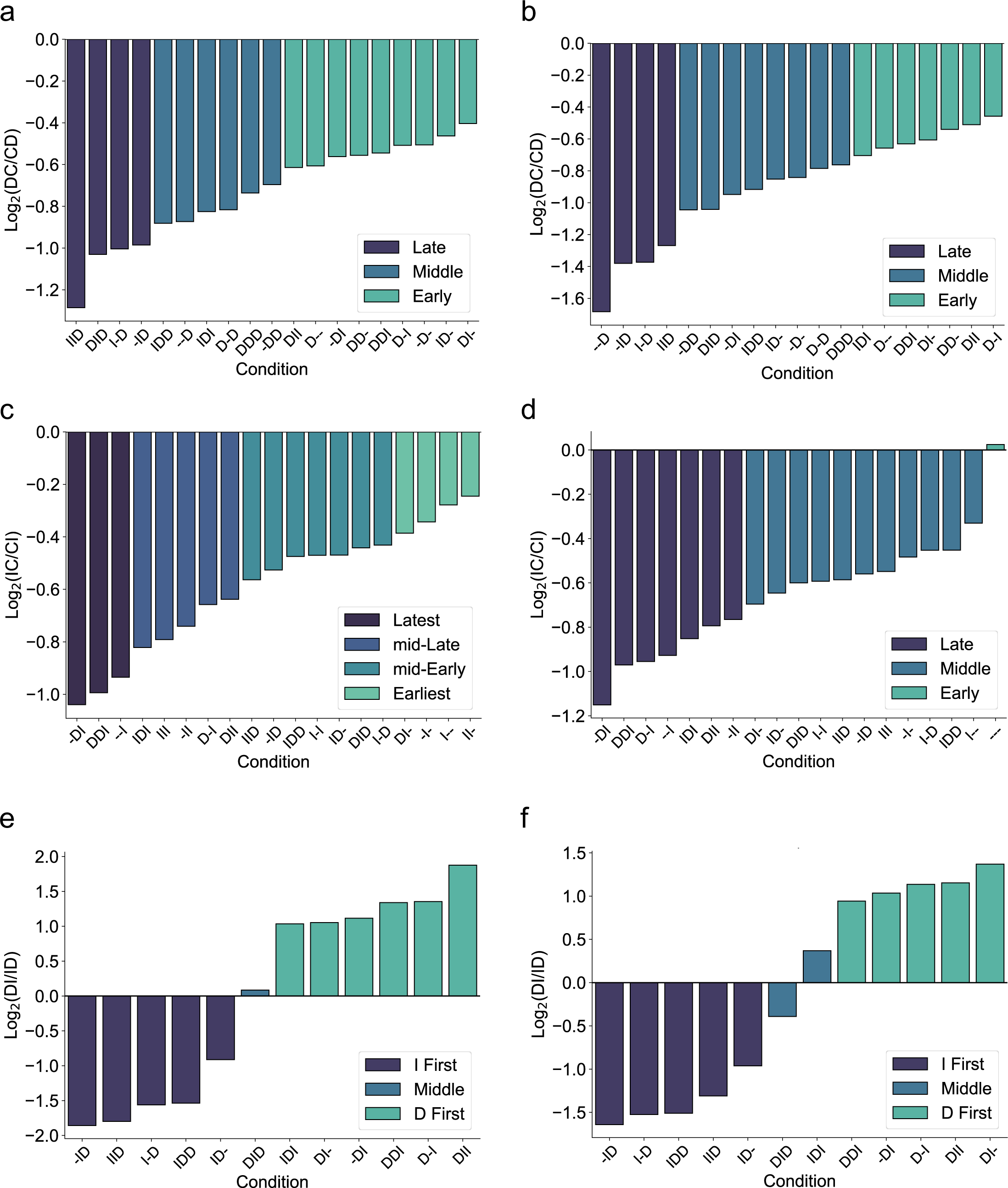
Recording of inducer order with peCHYRON. **a**, As described for Figure 5b, ratios of DC to CD bigrams were calculated. In this case, we calculated bigrams only for those samples that did not cluster with the lowest D proportions in Extended Data Figure 8a. We then applied k-means clustering to the log_2_ of the bigram ratios, choosing the cluster number automatically as in Extended Data Figure 8, and colored bars based on their cluster membership. These data are for Replicate 1. **b**, As in a, for Replicate 2. **c**, As in a, but ratios of IC to CI bigrams. **d**, As in c, for Replicate 2. **e**, As in a, but ratios of DI to ID bigrams. **f**, As in e, for Replicate 2.

**Extended Data Figure 10.**
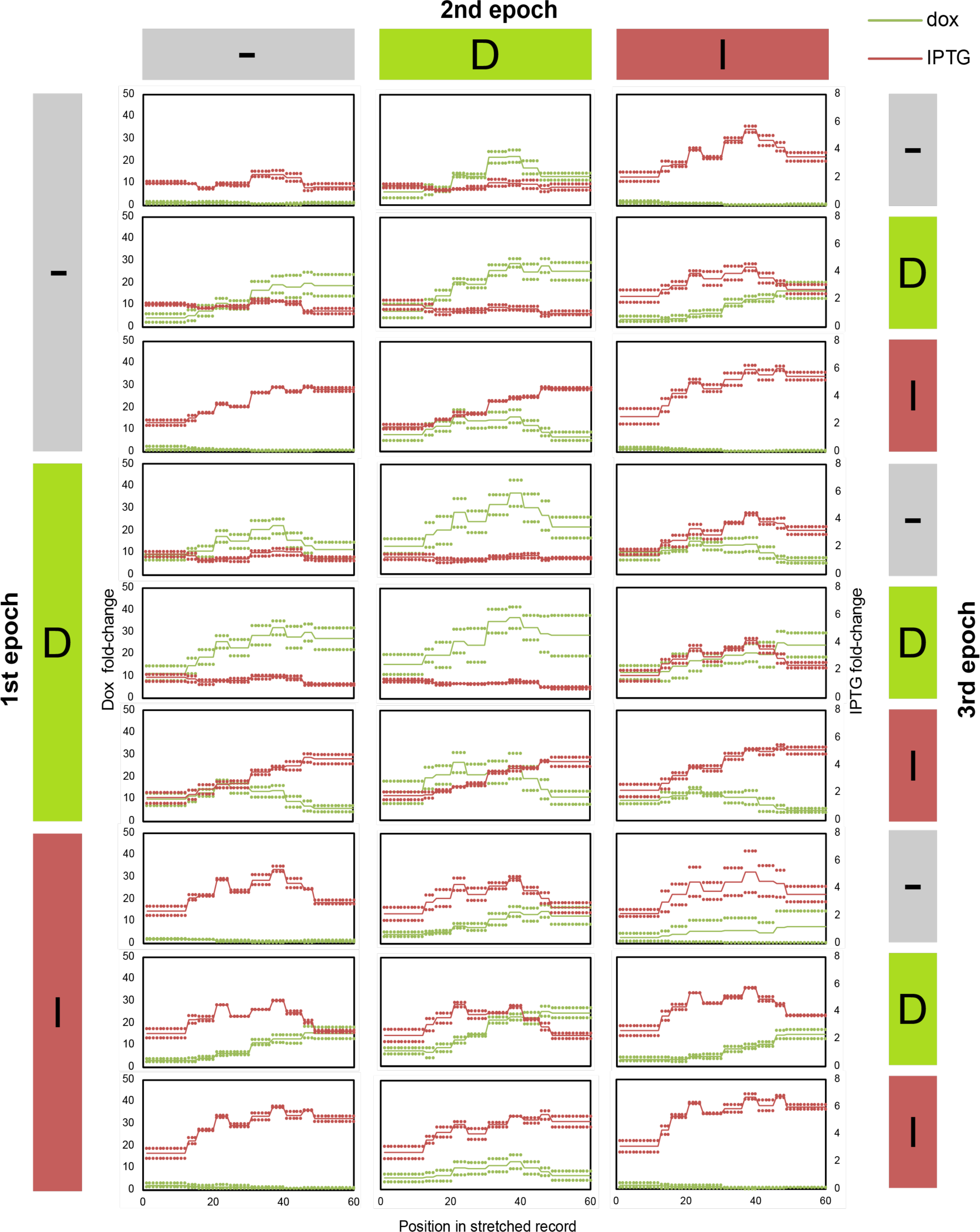
Signature location on the peCHYRON array as a proxy for time. Stretched records underlying the analysis shown in Figure 5c. Points represent biological replicates and lines are the mean of the biological replicates. Data is organized according to the inducer, or lack thereof, that each sample was exposed to during the 3 epochs.

