## Supplementary Table 3 for "Open-ended molecular recording of sequential cellular events into DNA"

|  | peCHYRON | DNA Typewriter |
| --- | --- | --- |
| Number of components needed | 4: prime editor, protective nicking sgRNA, A→B pegRNA, B→A pegRNA | 3: prime editor, pegRNA, synthetic recording locus |
| Recording loci | Can be synthetic or natural genomic sites. | Synthetic loci must be integrated. |
| Locus length per record | 20 bp | 18.4 bp |
| Compatibility with enhanced pegRNAs with protected 3’ ends | Not in its current form. | Yes. |
